# Coro1A and TRIM67 collaborate in netrin-dependent neuronal morphogenesis

**DOI:** 10.1101/2025.03.20.644333

**Authors:** Chris T. Ho, Elliot B. Evans, Kimberly Lukasik, Ellen C. O’Shaughnessy, Aneri Shah, Chih-Hsuan Hsu, Brenda Temple, James E. Bear, Stephanie L. Gupton

## Abstract

Neuronal morphogenesis depends on extracellular guidance cues accurately instructing intracellular cytoskeletal remodeling. Here, we describe a novel role for the actin binding protein Coronin 1A (Coro1A) in neuronal morphogenesis, where it mediates responses to the axon guidance cue netrin-1. We found that Coro1A localizes to growth cones and filopodial structures and is required for netrin-dependent axon turning, branching, and corpus callosum development. We previously discovered that Coro1A interacts with TRIM67, a brain enriched E3 ubiquitin ligase that interacts with a netrin receptor and is also required for netrin-mediated neuronal morphogenesis. Loss of Coro1A and loss of TRIM67 shared similar phenotypes, suggesting that they may function together in the same netrin pathway. A Coro1A mutant deficient in binding TRIM67 was not able to rescue loss of Coro1A phenotypes, indicating that the interaction between Coro1A and TRIM67 is required for netrin responses. Together, our findings reveal that Coro1A is required for proper neuronal morphogenesis, where it collaborates with TRIM67 downstream of netrin.

## Introduction

During brain development, neurons undergo rapid shape changes that are key to establishing functional neural circuitry. Integral to appropriate morphogenesis are the extracellular cues that inform directional growth via spatiotemporal remodeling of cytoskeletal networks (Kolodkin and Tessier-Lavigne, 2011; Lowery and Vactor, 2009; Dent et al., 2011; McCormick and Gupton, 2020; Ho and Gupton, 2021). The secreted guidance cue netrin-1 (netrin) plays a critical role during axon pathfinding. In mice, loss of netrin or its receptor DCC results in axon guidance defects (Serafini et al., 1996; Fazeli et al., 1997; Bin et al., 2015; Yung et al., 2015), including agenesis of the corpus callosum and hippocampal commissure. Beyond its classic guidance role, netrin also promotes axon branching and synaptogenesis during development (Dent et al., 2004; Flores, 2011; Goldman et al., 2013; Glasgow et al., 2018; Wong et al., 2019). Numerous works have shown that netrin-dependent neuronal shape changes require the reorganization of the actin cytoskeleton (Boyer and Gupton, 2018). However, our understanding of how netrin interfaces with diverse intracellular actin cytoskeleton machinery to propagate myriad responses during neuronal morphogenesis remains incomplete.

The tripartite motif (TRIM) family of RING-domain containing E3 ubiquitin ligases play critical roles during neuronal development (Tocchini and Ciosk, 2015; Dudley-Fraser and Rittinger, 2023), in particular class-I TRIM proteins. Phylogenetic analysis suggest that class-I TRIMs emerged alongside the evolution of neurons (Boyer et al., 2018). Vertebrates have six class-I TRIMs, whereas invertebrates have one or two. The TRIM67 and TRIM9 paralogs are the most evolutionary conserved class-I TRIM and are enriched in the brain during development (Short and Cox, 2006; Boyer et al., 2018). In *Drosophila* and *C. elegans*, the loss of the single class-I TRIM (*MADD-2*/*Trim9*) results in axon guidance and branching defects (Hao et al., 2010; Morikawa et al., 2011), phenocopying loss of the invertebrate ortholog of *Ntn1* or *Dcc*. In mice, TRIM67 and TRIM9 interact with DCC and are independently required for netrin-dependent neuronal morphogenesis including growth cone morphology, axon turning, axon branching, and synaptogenesis (Winkle et al., 2014; Menon et al., 2015; Boyer et al., 2018; Winkle et al., 2016; Plooster et al., 2017; Boyer et al., 2020; McCormick et al., 2024a; Mutalik et al., 2025). However, unlike invertebrates, loss of either TRIM67 or TRIM9 does not mimic the phenotypes observed with loss of netrin or DCC. Furthermore, *Trim67*^-/-^ and *Trim9*^-/-^ mice display opposing phenotypes. For example, *Trim67*^-/-^ mice exhibit a thinner corpus callosum (Boyer et al., 2018, 2020), whereas *Trim9*^-/-^ have a thicker corpus callosum (Winkle et al., 2014; Menon et al., 2015; Plooster et al., 2017). These findings suggest that TRIM67 and TRIM9 are independently required downstream of netrin and DCC.

E3 ubiquitin ligases typically have multiple substrates and interaction partners. Using unbiased proximity-dependent biotin identification (BioID), we identified shared and unique TRIM9 and TRIM67 binding partners in developing cortical neurons (Menon et al., 2021). The actin binding protein coronin 1A (Coro1A) was identified as an interactor of TRIM67, but not TRIM9. Coro1A is a member of the coronin family. Coronin was originally discovered in *Dictostelium* (de Hostos et al., 1991), where it localized to the “crown-like” F-actin protrusions on the dorsal cell surface. Coronins are characterized as essential actin binding proteins, conserved across a wide range of eukaryotes, from yeast to human (Chan et al., 2011). Coro1A along with three other coronins belong to the type I coronin class. Type I coronins are the most studied and closely resemble ancient coronins. Type I coronins play fundamental roles in diverse processes (Chan et al., 2011), including cell migration and membrane trafficking (Cai et al., 2005; Föger et al., 2006; Cai et al., 2007a; b; Hoyer et al., 2018; Striepen and Voeltz, 2022; King et al., 2022). Coronins regulate actin dynamics via modulating the Arp2/3 actin branching complex and ADF/cofilin (hereafter referred to as cofilin) actin severing proteins. Coronins interact with the Arp2/3 complex and inhibit Arp2/3 nucleating activity (Humphries et al., 2002; Rodal et al., 2005; Cai et al., 2007b). In addition, coronins promote debranching of Arp2/3-nucleated branched actin network (Cai et al., 2008). Coronin and cofilin cooperate in actin depolymerization (Goode et al., 1999; Brieher et al., 2006; Cai et al., 2007b). In mammalian cells, coronin promotes cofilin activity via recruiting its activator Slingshot phosphatase (SSH1L) (Cai et al., 2007b). Type I coronins primarily localize to the lamellipodia, where actin forms an intricate branched network. Interestingly, type I coronin members have distinct expression patterns (Nal et al., 2004; Cai et al., 2005; Chen et al., 2014; Spoerl et al., 2002). Coronin 1B and Coronin 1C are ubiquitously expressed in all mammalian cell types (Uetrecht and Bear, 2006; Chan et al., 2011). Coro1A is primarily expressed in hematopoietic cells and to a lesser extent in neurons. In sympathetic neurons, Coro1A is required for nerve growth factor and TrkA receptor signaling (Suo et al., 2014, 2015), a critical signaling pathway for peripheral nervous system development (Bodmer and Kuruvilla, 2012). In excitatory cortical neurons, Coro1A regulates the excitatory-to-inhibitory synapse ratio (Jayachandran et al., 2014). Despite these findings, our understanding of the functions of Coro1A during neuronal morphogenesis is still limited, particularly in early cortical neuron development. Given the interaction between Coro1A and TRIM67, and the role of coronins in regulating actin turnover and architecture, we hypothesized that Coro1A is involved in TRIM67-mediated neuronal shape changes and responses to netrin.

Here we uncovered a novel role for Coro1A in neuronal morphogenesis. We demonstrate that Coro1A protein increases during cortical neuron maturation in vitro and in vivo. Coro1A is enriched in growth cones of developing neurons and localizes to the base of filopodia structures. Loss of Coro1A shares similar phenotypes to loss of TRIM67, including increased growth cone size and deficits in netrin-mediated axon guidance as well as branching in vitro, and a reduction in the size of the developing corpus callosum. Surprisingly, we found this to be a ubiquitin independent regulation. We identified the domains essential for the Coro1A:TRIM67 interaction and demonstrated that the interaction between Coro1A and TRIM67 is key to neuronal responses to netrin. Together, these findings indicate that Coro1A and TRIM67 collaborate in netrin-dependent morphogenesis in cortical neurons.

## Results

### Coro1A is a TRIM67 interacting partner

We recently identified Coro1A as a TRIM67 binding partner in developing cortical neurons using BioID, a proximity-labeling mass spectrometry technique (Menon et al., 2021). The mammalian genome comprises seven coronins classified into type I, II, and III based on their protein structure and sequence similarity (Uetrecht and Bear, 2006; Chan et al., 2011). Coro1A is a type I coronin along with Coronin 1B (Coro1B), Coronin1C (Coro1C) and Coronin 6 (Coro6) (Fig. 1A). Multiple rodent brain RNAseq datasets show that Coro1A, Coro1B, and Coro1C are expressed in developing neurons, whereas little to no Coro6 transcript was detected (Fig S1A) (Zhang et al., 2014; Loo et al., 2019). We set out to determine whether neuronally expressed type I coronins interact with TRIM67, or whether the interaction was unique to Coro1A, as suggested by BioID. To do so, myc-tagged full length TRIM67 and GFP-tagged type I coronins were expressed in human embryonic kidney (HEK) 293 cells. As anticipated, myc-TRIM67 coimmunoprecipitated Coro1A-GFP (Fig. 1B). In contrast, little to no myc-TRIM67 was detected when Coro1B-GFP or Coro1C-GFP were immunoprecipitated. Notably, at least five times more myc-TRIM67 coimmunoprecipitated with Coro1A-GFP than with either Coro1B-GFP or Coro1C-GFP in all three biological replicates. This indicates TRIM67 specifically interacts with Coro1A. To confirm the interaction occurred in neurons, as suggested by BioID, we performed coimmunoprecipitation assays in primary cultured cortical neurons. Using Coro1A or TRIM67 antibodies, we were unable to detect an interaction between endogenous TRIM67 and Coro1A by coimmunoprecipitation. However, interactions between E3 ubiquitin ligases and binding partners are frequently transient and difficult to detect (Kim et al., 2015; Iconomou and Saunders, 2016), and indeed motivated our BioID study. As an alternative we increased Coro1A levels by transducing neurons with Coro1A-GFP adenovirus and used high affinity GFP-TRAP beads to increase the efficiency of Coro1A precipitation. Endogenous TRIM67 was present in the precipitate with Coro1A-GFP (Fig. 1C), but not in the non-transduced precipitate. These data confirm that Coro1A and TRIM67 interact in neurons.

**Figure 1.**
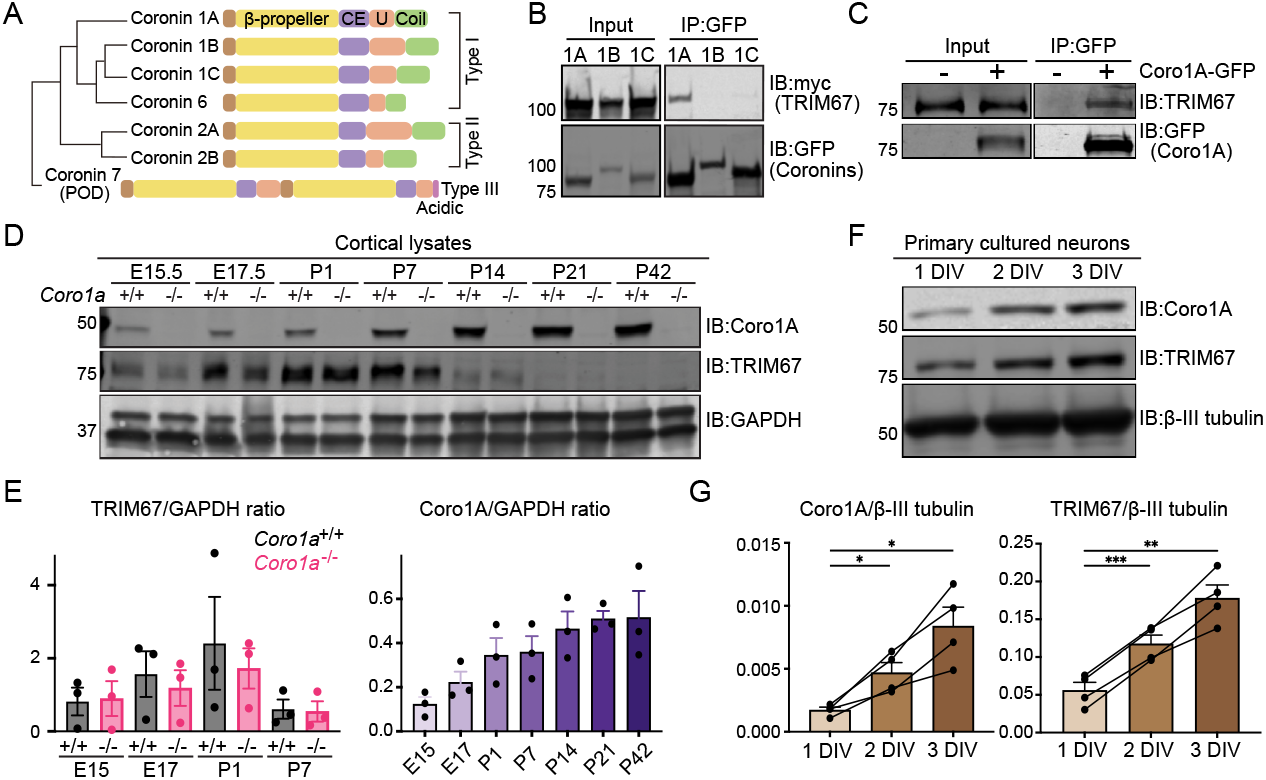
Coro1A interacts with TRIM67 and its protein level increases across neuron development. (**A**) Phylogenetic genetic tree of mammalian coronin family adapted from (Uetrecht and Bear, 2006; Chen et al., 2014). (**B**) Coimmunoprecipitation assays from HEK293 cells transfected with myc-TRIM67 and brain expressed coronins tagged with GFP demonstrating TRIM67 specifically interacts with Coro1A, but not Coro1B and Coro1C. Western blot is representative of three biological repeats. (**C**) Coimmunoprecipitation assays from 3 DIV cultured cortical neurons transduced with Coro1A-GFP adenovirus or non-transduced demonstrating that endogenous TRIM67 coimmunoprecipitates with Coro1A-GFP. Western blot is representative of two biological repeats. (**D**) Western blotting of *Coro1a*^+/+^ and *Coro1a*^-/-^ cortical lysate from indicated embryonic (E) and postnatal (P) ages showing Coro1A and TRIM67 expression patterns. (**E**) Quantification of TRIM67 and Coro1A protein level in cortical lysates normalized to GAPDH. n (animals) =3 per condition. Error bars denote SEM. (**F**) Western blotting of cultured *Coro1a*^+/+^ cortical neurons collected at 1 DIV, 2 DIV and 3 DIV showing increased levels of Coro1A and TRIM67 across neuron development. (**G**) Quantification of TRIM67 and Coro1A protein level in cultured neurons normalized to β-III-tubulin. n(cultures)= 4. Error bars denote SEM. *, p<0.05; **, p<0.01; ***, p<0.001. One-way ANOVA (Bonferroni’s post hoc test).

### Coro1A protein increases during neuronal development

We next investigated Coro1A and TRIM67 expression during cortical neuron development. We collected cortices from *Coro1a*^+/+^ and *Coro1a*^-/-^ littermates at developmental timepoints spanning stages of axon/dendrite morphogenesis and dendritic spine formation and synapse maturation (Embryonic day (E)15, E17, postnatal day (P)1, P7, P14, P21 and P42). Coro1A protein increased across cortical development (Fig. 1D-E). TRIM67 protein increased during embryonic stages, peaked at late embryonic and early postnatal time points, and subsequently decreased in juvenile and adolescent brains (Fig. 1D-E) as described (Boyer et al., 2018). Coro1A is enriched in microglia cells (Fig S1B)(Ahmed et al., 2007), which proliferate throughout early postnatal development (Nikodemova et al., 2015; Thion and Garel, 2017; Askew and Gomez-Nicola, 2018; Wurm et al., 2021). Thus, the increased Coro1A protein level may be due to an increase of microglia. To confirm that Coro1A was present in neurons during morphogenesis, we used cultured cortical neurons. Coro1A was detected at 1 DIV, and its expression level further increased at 2 and 3 DIV (Fig. 1F-G). A similar pattern was observed for TRIM67 (Fig. 1F-G). We demonstrated that TRIM67 was a component of the postsynaptic density (PSD) (Mccormick et al., 2024). Previous studies indicated that Coro1A was enriched in excitatory synapses (Jayachandran et al., 2014). Using subcellular fractionation of P21 mouse forebrains to enrich for the PSD fraction, we identified Coro1A in the PSD (Fig S1C). These data suggest that TRIM67 and Coro1A are present in developing cortical neurons.

### Coro1A is enriched in neuronal growth cones at the base of the filopodia

Type 1 coronins bind to filamentous actin (F-actin) and are enriched at the leading edge of migrating cells (Mishima and Nishida, 1999; Cai et al., 2005; King et al., 2022). We hypothesized that Coro1A may localize to the neuronal growth cone, which has a similar actin architecture to the leading edge (Svitkina and Borisy, 1999; Svitkina et al., 2003; Pollard and Cooper, 2009), including branched actin filament networks in lamellipodial veils and prominent bundled actin within filopodial protrusions (Lewis and Bridgman, 1992; Korobova and Svitkina, 2008). To assess Coro1A localization, we paraformaldehyde fixed cortical neurons at 2 DIV and stained for Coro1A, F-actin, and βIII-tubulin. Coro1A localized to F-actin rich growth cone structures and appeared as diffuse puncta in the soma (Fig. 2A). Growth cones are dynamic structures at the tip of a growing axon/immature neurite (Dent and Gertler, 2003; Lowery and Vactor, 2009). Because the dynamic remodeling of the F-actin is integral to the growth cone, we were interested in the dynamics of Coro1A in the growth cone. We expressed Coro1A-GFP in cortical neurons at 2 DIV when the growth cone is actively extending immature neurites. Total internal reflection fluorescence (TIRF) time-lapse microscopy revealed prominent localization of Coro1A-GFP in the growth cones that underwent retrograde flow (Fig. 2B and Supplemental Video 1,2). Coro1A-GFP localized to growth cone filopodial protrusions. This localization to bundled actin was surprising, as type I coronins are classically described as branched actin regulators.

**Figure 2.**
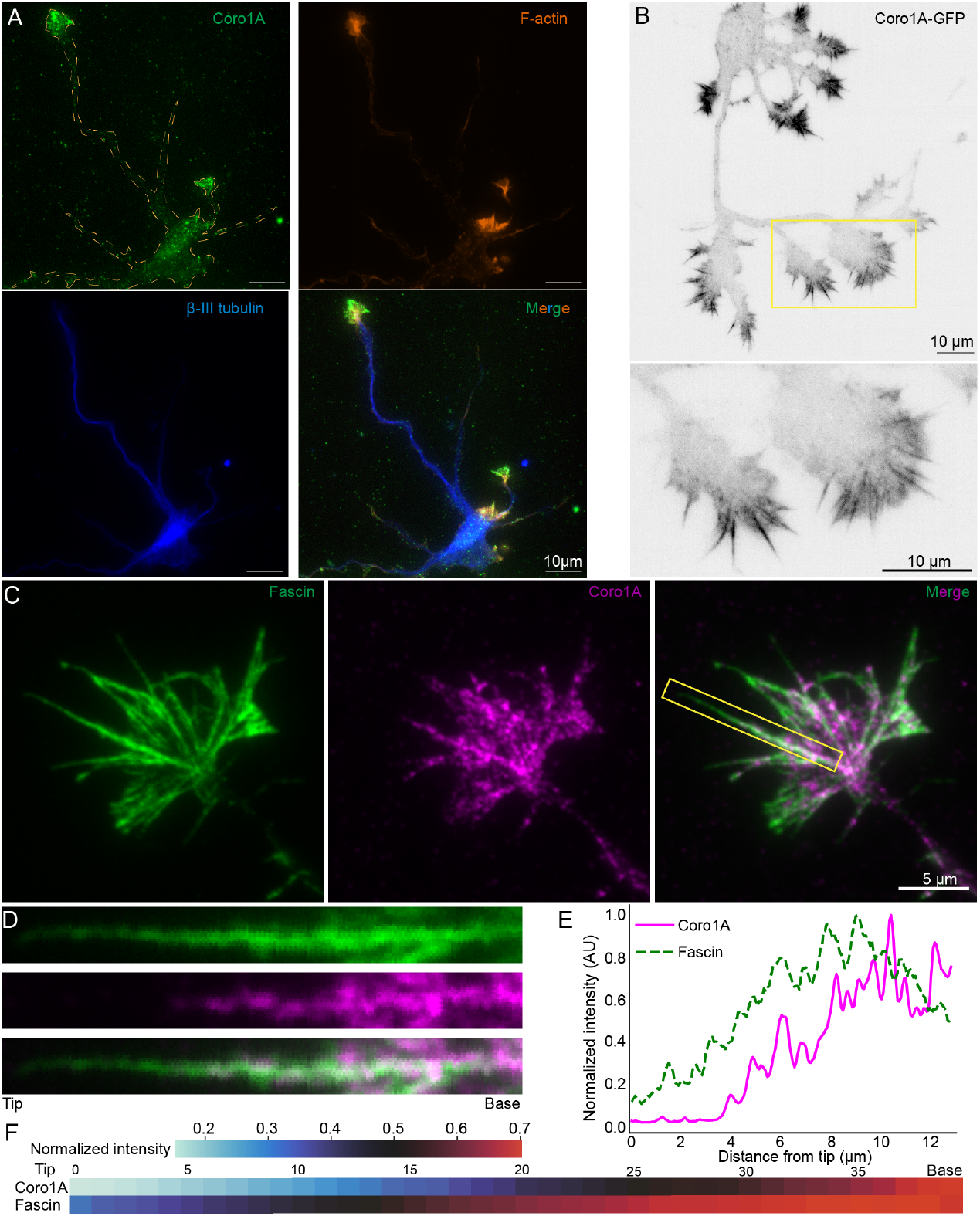
Coro1A localizes to neuronal growth cones and filopodia. (**A**) Representative images of 2 DIV cortical neurons fixed with paraformaldehyde and stained for Coro1A, β-III-tubulin, and filamentous actin (phalloidin) to demonstrate that endogenous Coro1A localizes to growth cones. (**B**) The top panel is a representative image from total internal reflection fluorescence (TIRF) live-cell movie of 2 DIV cortical neuron expressing Coro1A-GFP. Coro1A localizes to growth cones and growth cone filopodia protrusions. The bottom panel is a zoomed-in image from the yellow box in the top panel. (**C**) Representative images of 2 DIV cortical neuron fixed with ice-cold methanol and stained for fascin and Coro1A to demonstrate Coro1A localizing to the growth cone filopodia. (**D**) Inset of images from yellow box in C, showing that Coro1A is enriched at the base of the filopodia. (**E**) Representative line scan intensity profile of a single filopodium, filopodium shown in D, showing Coro1A and fascin distribution. Line scan was drawn from the tip to the base of the filopodium. The fluorescent intensity of each protein was normalized to the maximum intensity measured across the filopodium. (**F**) Heatmaps displaying the sub-filopodial localization of Coro1A and fascin averaged across n=706 filopodia from three biological replicates.

To confirm that the localization of Coro1A-GFP in filopodia was not an artifact of overexpression or tagging, we adapted our fixation method to ice-cold methanol, which enhanced Coro1A endogenous immunostaining. This enhanced staining phenomenon was previously observed for cofilactin in neurons (Hylton et al., 2022; Minamide et al., 2023). We immunostained 2 DIV neurons for Coro1A and fascin, an actin binding protein that bundles actin within the filopodia. We observed Coro1A on fascin-labeled growth cone filopodia (Fig. 2C). We stained *Coro1a*^+/+^ and *Coro1a*^-/-^ cultured neurons to validate antibody specificity (Fig. S2). We examined Coro1A sub-filopodia localization intensity distribution across multiple filopodia. Fascin staining runs from filopodium tip to the base, whereas Coro1A staining began in the middle region of the filopodium shaft and extended to the base (Fig. 2D-E). We quantified 706 filopodia demonstrating that Coro1A was enriched at the base (the distal 50%) (Fig. 2F). Taken together, these findings show that Coro1A localizes to growth cones and is enriched in growth cone filopodia, particularly at the base region in developing neurons.

### Coro1A mediates netrin-dependent axon guidance

Appropriate axon guidance depends on response to extracellular guidance cues and subsequent remodeling of the growth cone cytoskeleton. Coro1A is enriched in the growth cone and interacts with TRIM67, a brain-enriched E3 ubiquitin ligase that functions downstream of netrin (Boyer et al., 2018, 2020). We hypothesized that Coro1A may function in netrin-dependent axon guidance. To test this, we performed in vitro axon turning assays with *Coro1a*^+/+^ and *Coro1a*^-/-^ cortical neurons, utilizing a microfluidic axon guidance chamber developed in our lab (Taylor et al., 2015). These chambers enable fluidic isolation of axons from the somatodendritic compartment and establishment of a stable netrin gradient within the region only accessible to axons (Fig. 3A). A dextran gradient serves as a negative control to ensure that fluidic flow does not induce turning (Supplemental Video 3). We measured the angle between the initial and final orientation of the axon (Fig. 3B). A positive turning angle indicates attractive turning toward a netrin source, whereas a negative angle represents repulsive turning. Wildtype neurons exhibited a biphasic response in a netrin gradient (Supplemental Video 4-5), as reported (Taylor et al., 2015; Boyer et al., 2020; Mutalik et al., 2025). At the low end of the netrin gradient, *Coro1a*^+/+^ axons showed positive turning angles (attractive response). At the high end of the gradient *Coro1a*^+/+^ axons exhibited negative turning angles (repulsive turning). *Coro1a*^-/-^ axons failed to show turning responses to netrin at either end of the gradient (Fig. 3C, Supplemental Video 4-5). Rather they exhibited small and random turning angles, similar to the dextran-only gradient. These axon turning assays demonstrate that Coro1A is required for netrin-mediated axon guidance.

**Figure 3.**
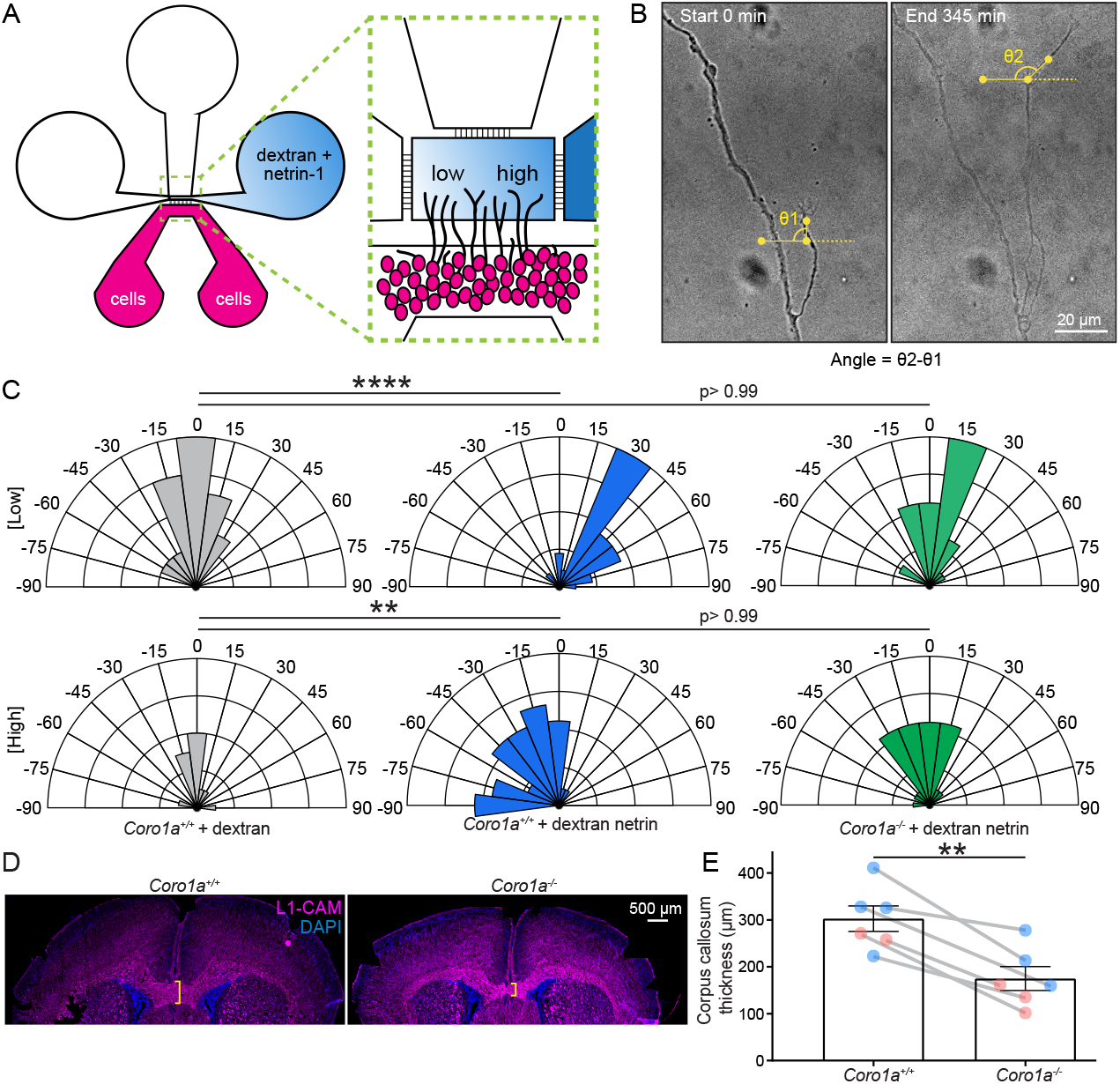
Coro1A is required for netrin-dependent axon guidance in vitro and in vivo. (**A**) Schematic of microfluidic device for axon turning assays. (**B**) Example of a *Coro1a*^+/+^ axon that entered axon viewing area and turned toward the high netrin. Analysis procedure for defining axon turning angle. (**C**) Quantification of axon turning angle shown by rose plots. Angles > 90° or < -90° were grouped into 90° and -90°, respectively. n(cells)= 29 *Coro1a*^+/+^ dextran low, 24 *Coro1a*^+/+^ netrin low, 25 *Coro1a*^-/-^ netrin low, 11 *Coro1a*^+/+^ dextran high, 35 *Coro1a*^+/+^ netrin high, 25 *Coro1a*^-/-^ netrin high from four biological replicates. One-way ANOVA (Bonferroni’s post hoc test). **, p<0.01; ****, p<0.0001. (**D**) Representative coronal sections of the P4 brains from *Coro1a*^+/+^ and *Coro1a*^-/-^ littermates stained with L1-CAM and DAPI. (**E**) Measurement of L1CAM^+^ corpus callosum thickness. n (animals)= 6 *Coro1a*^+/+^ (4 males, 2 females) and 6 *Coro1a*^-/-^ (3 males, 3 females). Paired two-tailed t-test. **, p<0.01.

Deletion of *Ntn1* or its receptor *Dcc* results in agenesis of the corpus callosum (Serafini et al., 1996; Fazeli et al., 1997). *Trim67*^-/-^ mice have a thinner corpus callosum (Boyer et al., 2018, 2020). We next assessed this netrin-sensitive axon tract in *Coro1a*^-/-^ mice. We performed serial coronal sections of P4 *Coro1a*^+/+^ (four males and two females) and *Coro1a*^-/-^ (three males and three females) littermate paired brains and measured the size of the corpus callosum at its most rostral position (Fig. 3D). This position corresponds to first anterior section where the callosum appears, making comparison consistent between paired samples across litters. In all six pairs, the corpus callosum was thinner in the *Coro1a*^-/-^ mice compared to *Coro1a*^+/+^ littermate control (Fig. 3E). Taken together, Coro1A is involved in netrin-dependent axon guidance both in vitro and in vivo.

### Coro1A mediates netrin-dependent growth cone morphology and axon branching

Given that loss of Coro1A results in loss of turning responses in vitro and disruption of a netrin-sensitive projection in vivo, we next investigated whether Coro1A plays a role in neuronal growth cone morphology in response to netrin. Acute addition of netrin promoted expansion of the growth cone area in *Coro1a*^+/+^ neurons (Fig. 4A-B), in agreement with previous reports (Menon et al., 2015; Boyer et al., 2020; Mutalik et al., 2025). However, netrin treatment did not increase the area of the *Coro1a*^-/-^ growth cones, indicating Coro1A is required for growth cone response to netrin. Interestingly, at baseline prior to addition of netrin, *Coro1a*^-/-^ growth cones were larger than *Coro1a*^+/+^ (Fig. 4A-B). These phenotypes were similar to those exhibited by *Trim67*^-/-^ neurons (Boyer et al., 2020), suggesting that Coro1A potentially functions with TRIM67.

**Figure 4.**
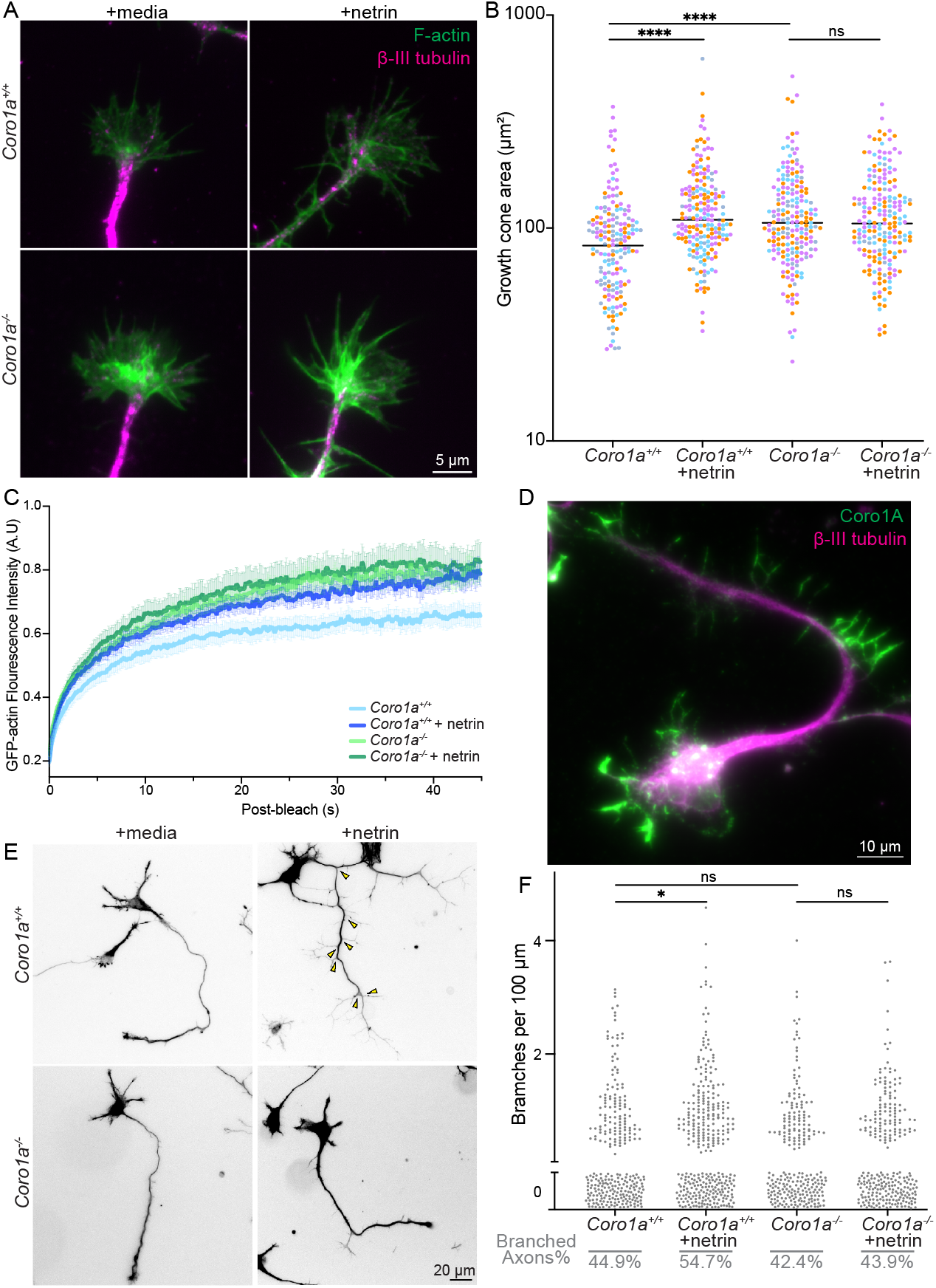
Coro1A is required for netrin-dependent growth cone morphology and axon branching. (**A**) Representative images of 2 DIV axonal growth cones from E15.5 *Coro1a*^+/+^ and *Coro1a*^-/-^ littermates treated with media or netrin (600ng/mL) for 40 min. Neurons were stained for β-III-tubulin and filamentous actin (phalloidin). (**B**) Quantification of growth cone area. n(cells)= 178 *Coro1a*^+/+^ media, 204 *Coro1a*^+/+^ netrin, 199 *Coro1a*^-/-^ media, 204 *Coro1a*^-/-^ netrin from three biological replicates. One-way ANOVA (Bonferroni’s post hoc test). ***, p < 0.001; ns, not statistically different. (**C**) Normalized fluorescence intensity of GFP-actin after photobleaching. The lines are mean normalized intensity, and the error bars are ± SEM. n(cells)= 15 *Coro1a*^+/+^, 20 *Coro1a*^+/+^ netrin, 20 *Coro1a*^-/-^ media, 13 *Coro1a*^-/-^ netrin from three biological replicates. (**D**) Representative image of 3 DIV neuron stained for β-III-tubulin and Coro1A, showing that Coro1A localizes to axonal filopodia. (**E**) Representative images of 3 DIV neurons from E15.5 *Coro1a*^+/+^ and *Coro1a*^-/-^ littermates treated with media or netrin (250 ng/mL) for 24h. Images were shown as the combined staining of β-III-tubulin and filamentous actin (phalloidin). Yellow arrowhead denotes branches. (**F**) Quantification of axonal branches per 100 µm axon length. n(cells)= 289 *Coro1a*^+/+^ media, 339 *Coro1a*^+/+^ netrin, 257 *Coro1a*^-/-^ media, 273 *Coro1a*^-/-^ netrin from three biological replicates. Kruskal-Wallis nonparametric ANOVA test (Dunn’s post hoc test). *, p <0.05; ns, not statistically different.

Since the growth cone is enriched in dynamic actin, the expansion of growth size in *Coro1a*^-/-^ neurons prompted us to investigate how loss of Coro1A affects actin turnover. We performed fluorescence recovery after photobleaching (FRAP) assays in the growth cones of neurons expressing GFP-actin (Fig. 4C, Supplemental Video 6-9). Acute netrin addition led to an increase in the percentage of fluorescent recovery of GFP-actin (mobile fraction) in *Coro1a*^+/+^ neurons (Fig.4C, S3A-B). This suggests that netrin increases actin turnover, decreasing the immobile fraction of actin in the growth cone. Consistent with the growth cone morphology results, the percentage of GFP-actin fluorescent recovery in *Coro1a*^-/-^ neurons was unchanged when treated with netrin (Fig. 4C, S3A-B), suggesting that Coro1A is required for actin responses to netrin in the growth cone. We found no difference in fluorescence recovery halftime following netrin treatments in both genotypes (Fig S3C), indicating a similar rate of actin turnover and/or retrograde flow.

Netrin also promotes axon branching (Dent et al., 2004; Winkle et al., 2014; Boyer and Gupton, 2018; Boyer et al., 2020). We observed that Coro1A localizes to filopodia along the axons (Fig. 4D), which are considered axon branch precursors (Bastmeyer and O’Leary, 1996; Spillane et al., 2011; Gallo, 2011; Kalil and Dent, 2014; Ho and Gupton, 2021). Thus, we examined whether netrin-dependent axon branching involves Coro1A. *Coro1a*^+/+^ neurons significantly increased axon branching density 24 h after addition of netrin (Fig. 4E-F). However, *Coro1a*^-/-^ neurons did not increase axon branching density. These data demonstrate that Coro1A is required for multiple netrin-dependent responses during neuronal morphogenesis.

### Coro1A multi-monoubiquitination independent of TRIM67 or netrin

Since Coro1A and TRIM67 interact, and *Trim67*^-/-^ and *Coro1a*^-/-^ neurons lose netrin-dependent responses during neuronal morphogenesis (Boyer et al., 2020), this prompted us to investigate whether Coro1A was ubiquitinated. We co-expressed Coro1A-GFP and hemagglutinin (HA)-ubiquitin in HEK293 cells. Following immunoprecipitation of Coro1A-GFP under denaturing conditions, we observed one HA-ubiquitin positive band that comigrated with Coro1A-GFP, and four discrete high-molecular-weighted ubiquitin bands (Fig. 5A). In contrast, none of these ubiquitin positive bands were present when immunoprecipitating GFP, suggesting that Coro1A is ubiquitinated. The discrete high-molecular-weight ubiquitin bands led us to suspect that Coro1A was monoubiquitinated or multi-monoubiquitinated, as we had observed similar discrete ubiquitin bands with the TRIM9/TRIM67 regulation of the actin polymerase VASP by multi-monoubiqutination (Menon et al., 2015; Boyer et al., 2020; McCormick et al., 2024b). To test this, we performed ubiquitination assays with HEK293 cells expressing Coro1A-GFP and HA-ubiquitin or HA-ubiquitin^K0^, a ubiquitin mutant incapable of forming ubiquitin chains. The discrete high molecular weight ubiquitin bands were present with both HA-ubiquitin or HA-ubiquitin^K0^ (Fig. 5B), suggesting that Coro1A is multi-monoubiquitinated. Finally, we explored whether this Coro1A ubiquitination was present in neurons. Unfortunately, we were unable to identify a coronin antibody compatible with immunoprecipitation under the denaturing conditions required and turned to adenoviral Coro1A-GFP transduction. After immunoprecipitating Coro1A-GFP from cultured neuronal lysates, we detected similar discrete HA-ubiquitin high-molecular-weighted positive bands (Fig. 5C), indicating that Coro1A is ubiquitinated in neurons. However, the ubiquitination was not altered by loss of *Trim67* or addition of netrin (Fig. 5D). Therefore, although Coro1A is likely multi-monoubiquitinated, this ubiquitination is independent of either TRIM67 or netrin.

**Figure 5.**
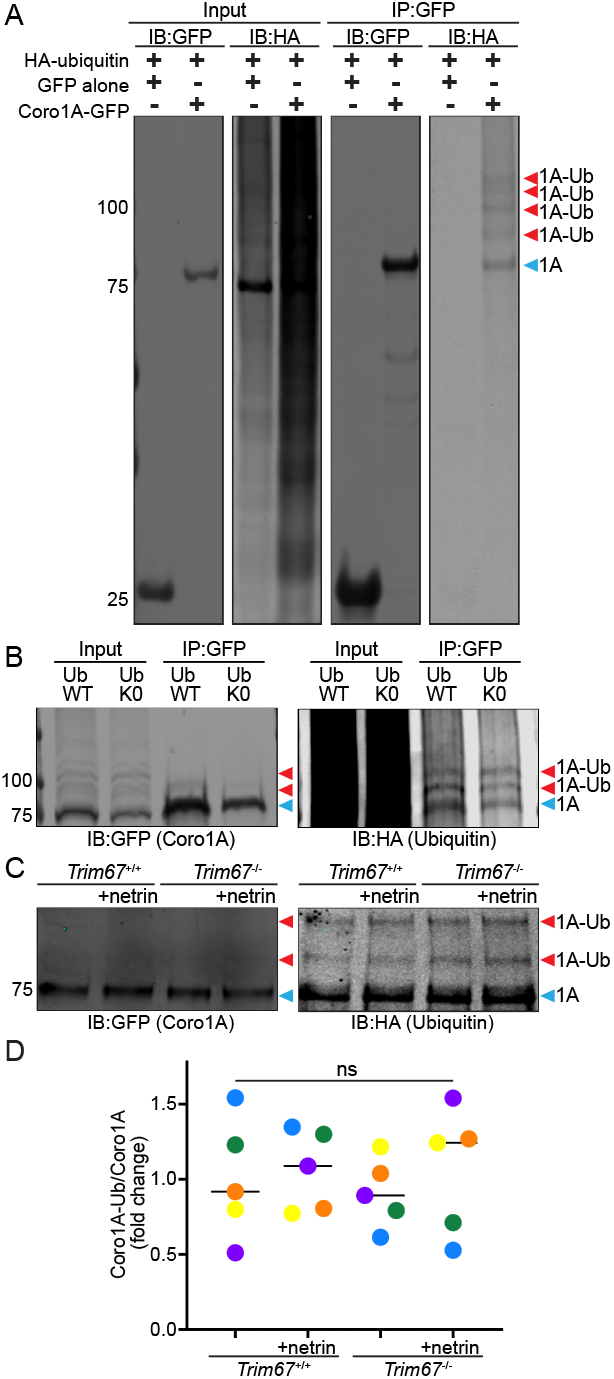
Coro1A monoubiquitination is independent of TRIM67 and netrin. (**A**) Western blot of Coro1A-GFP or GFP immunoprecipitated under denaturing conditions from HEK293 cells expressing HA-ubiquitin, showing Coro1A is ubiquitinated, as the higher-molecular weight HA^+^ bands (Coro1A-Ub, red arrowheads) were only present in the fraction precipitated by Coro1-GFP but not GFP. Western blot is representative of three biological repeats. (**B**) Western blot of Coro1A-GFP immunoprecipitated under denaturing conditions from HEK293 cells expressing HA-ubiquitin or HA-ubiquitin^K0^, demonstrating that Coro1A is monoubiquitinated. Western blot is representative of four biological repeats. (**C**) Western blot of Coro1A-GFP immunoprecipitated from denatured *Trim67* ^+/+^ or *Trim67*^-/-^ 3 DIV cortical neurons treated with or without netrin (600 ng/mL), demonstrating that Coro1A-Ub occurs in neurons, but is independent of either TRIM67 or netrin. (**D**) Western blot analysis of Coro1A-Ub normalized to total Coro1A level. n (cultures)= 5. One-way ANOVA (Bonferroni’s post hoc test). ns, not statistically different. Line indicated median for each condition.

### The coiled coil domain of Coro1A is essential for interaction with TRIM67

Since Coro1A ubiquitination was not altered by TRIM67, we next investigated whether their interaction was critical for netrin-dependent neuronal shape changes. To test this, we needed to generate mutants unable to interact for rescue experiments. This prompted us to define the domains required for binding.

TRIM67 (Fig. 6A), like other TRIM proteins, has an N-terminal tripartite motif that includes an E3 ubiquitin ligase RING domain (Meroni and Diez-Roux, 2005; Deshaies and Joazeiro, 2009), 2 BBox domains, and a coiled coil domain (Meroni and Diez-Roux, 2005). The C-terminus of TRIM67 consists of a Cos domain, FN3 domain, and SPRY domain (Short and Cox, 2006). Interactions between E3 ubiquitin ligases and their binding partners are typically transient and rapid. Removing the RING stabilizes protein-protein interaction. We found that removal of the RING domain of TRIM67 (TRIM67ΔRING) enhanced the interaction between TRIM67 and Coro1A (Fig. 6B-C). Of note, the TRIM67ΔRING also coimmunoprecipitated Coro1B-GFP and Coro1C-GFP (Fig S4A), yet significantly less than Coro1A-GFP (Fig S4B), further supporting a preferential binding of TRIM67 to Coro1A (Fig 1B). We utilized TRIM67ΔRING to enhance the Coro1A:TRIM67 interaction, to more robustly detect reductions in the interaction.

**Figure 6.**
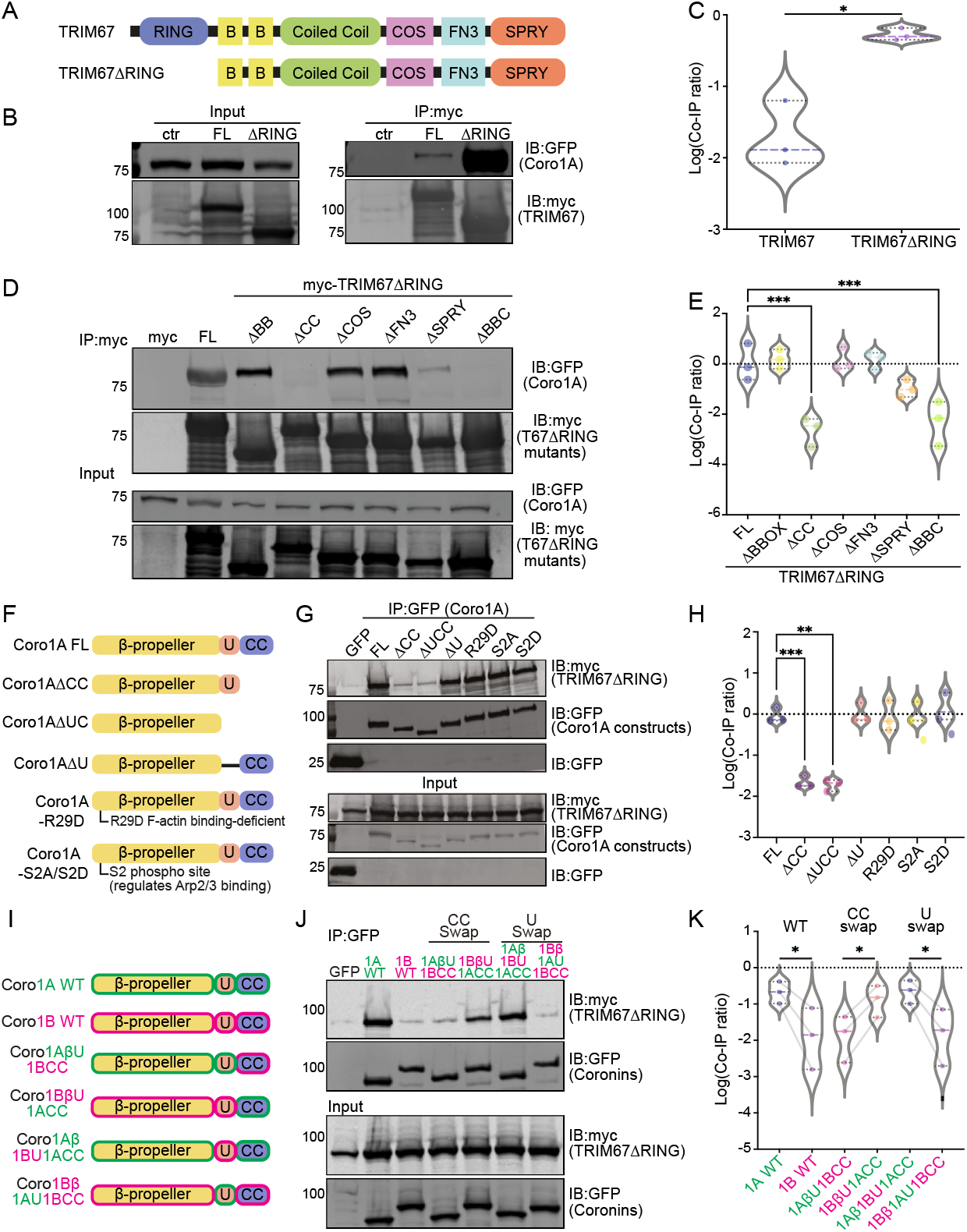
Coro1A coiled coil domain is required for Coro1A:TRIM67 interaction. (**A**) Schematic of TRIM67 domains and RING-truncated construct (TRIM67ΔRING). (**B**) Coimmunoprecipitation assays from HEK293 cells transfected with Coro1A-GFP and myc-TRIM67 or myc-TRIM67ΔRING demonstrating that deleting the RING domain enhances the interaction between Coro1A and TRIM67. (**C**) Quantification of Coro1A-GFP coimmunoprecipitation ratios (Co-IP ratios) from (B), showing Coro1A-GFP levels relative to myc-TRIM67 or myc-TRIMΔRING. n(cultures)= 3. Paired two-tailed t-test. *, p <0.05. (**D**) Structure-function coimmunoprecipitation assays from HEK293 cells transfected with Coro1A-GFP and myc-TRIM67ΔRING or indicated myc-TRIM67ΔRING domain deletion construct, showing that the coiled coil domain of TRIM67 is required for the Coro1A:TRIM67 interaction. (**E**) Quantification of Coro1A-GFP Co-IP ratios from (D), showing Coro1A-GFP levels relative to myc-TRIMΔRING constructs. n(cultures)= 3. One-way ANOVA (Dunnett’s multiple comparisons tests). ***, p <0.001. (**F**) Schematic of Coro1A domains and constructs used in structure-function assays. (**G**) Structure-function coimmunoprecipitation assays from *TRIM67*^-/-^ HEK293 cells transfected with myc-TRIMΔRING and indicated GFP-tagged Coro1A constructs, showing the requirement of Coro1A coiled coil domain for Coro1A:TRIM67 interaction. These assays also reveal the interaction is independent of Coro1A binding to either filamentous actin or the Arp2/3 complex. (**H**) Quantification of myc-TRIM67ΔRING Co-IP ratios from (G), showing myc-TRIM67ΔRING levels relative to Coro1A-GFP constructs. n(cultures)= 3. One-way ANOVA (Dunnett’s multiple comparisons tests). **, p <0.01; ***, p <0.001. (**I**) Schematic of coronin chimera constructs used to determine whether the coiled coil domain of Coro1A mediates Coro1A:TRIM67 interaction. (**J**) Structure-function CoIP assays from *TRIM67*^-/-^ HEK293 cells transfected with myc-TRIM67ΔRING and indicated GFP-tagged coronin constructs, showing Coro1A coiled coil domain mediates Coro1A:TRIM67 interaction. (**K**) Quantification of myc-TRIM67ΔRING Co-IP ratios from (J), showing myc-TRIM67ΔRING levels relative to the GFP tagged coronin chimera constructs. n(cultures)= 3. Paired one-tailed t-test. *, p <0.05.

We first mapped the domains of TRIM67 that were essential for the Coro1A:TRIM67 interaction using TRIM67ΔRING. Coro1A coprecipitated with most domain deletions, except for those lacking the coiled coil (Fig. 6D-E). Interestingly, we previously demonstrated that TRIM67-dependent regulation of netrin responses requires the coiled coil domain (Boyer et al., 2020). AlphaFold2 predictions suggested that amino acids 280-497 (BBox2-coiled coil-Cos, Fig S4C) is a long helix that forms an antiparallel dimer. This predicted dimerization region alone coimmunoprecipitated Coro1A (Fig S4D). Whereas the 159-279 (BBox1) and 489-768 region (FN3-SPRY) were unable to coimmunoprecipitate Coro1A (Fig S4D). Taken together, the interaction between Coro1A and TRIM67 required the coiled coil domain of TRIM67.

Next, we mapped the domains on Coro1A required for the Coro1A:TRIM67 interaction. Coro1A has three domains (Appleton et al., 2006). The NH2-terminus forms a large, seven-bladed β-propellor domain, considered to facilitate protein–protein interactions (McArdle and Hofmann, 2008; Stirnimann et al., 2010); the COOHterminus contains a coiled coil domain responsible for Coro1A trimerization (Kammerer et al., 2005; Gatfield et al., 2005; Liu et al., 2011; Han et al., 2023). These domains are separated by a unique region, which is intrinsically disordered and variable in sequence and length among coronins (Han et al., 2023)(Fig. 6F). Coro1A constructs lacking the coiled coil domain (Coro1AΔCC and Coro1AΔUCC) showed a significant reduction in TRIM67:Coro1A interaction. Deletion of the unique region alone (Coro1AΔU) did not affect the interaction between TRIM67:Coro1A (Fig. 6G-H). This suggested that the coiled coil domain, but not the unique region of Coro1A mediates strong binding between Coro1A and TRIM67.

However, the coiled coil domain mediates trimerization of coronins and is required for several protein functions (Goode et al., 1999; Asano et al., 2001; Humphries et al., 2002; Spoerl et al., 2002; Gandhi et al., 2009; Liu et al., 2011; Han et al., 2023), including actin binding, Arp2/3 inhibition, and cellular localization, complicating the interpretation of these binding experiments. In contrast to Coro1A, which displayed punctate and F-actin localization, the pattern of Coro1AΔCC appeared more diffuse (Fig S4E). Therefore, the reduction in interaction observed in the mutants lacking the coiled coil domain may have been due to loss of Coro1A trimerization. We were reluctant to perform binding assays with a mutant lacking the entire β-propeller that comprises 85% of the protein. We therefore targeted interface resides on the surface of the β-propeller. Since Coro1A has a stronger interaction with TRIM67 compared to Coro1B and Coro1C (Fig. 1B and S4A-B), we further predicted that the surface exposed residues on the β-propeller that were unique to Coro1A were integral for its interaction with TRIM67. To test this, we made a single construct with all 20 surface residues unique to Coro1A mutated to residues found on Coro1B or Coro1C (Fig S4F, Supplemental Table 1). Surprisingly, the mutated construct was able to coimmunoprecipitate an indistinguishable amount of TRIM67 to wildtype Coro1A (Fig S4G). These data suggested that the unique surface residues on Coro1A were not responsible for the Coro1A:TRIM67 interaction.

As Coro1A is an F-actin and Arp2/3 binding protein (Grogan et al., 1997; Machesky et al., 1997), we wondered whether the Coro1A:TRIM67 interaction required Coro1A binding to F-actin or the Arp2/3 complex. To investigate this, we took advantage of Coro1A mutants that do not interact with F-actin or the Arp2/3 complex. The surface-exposed conserved arginine (Arg29) in the β-propeller is required for Coro1A binding to F-actin (Cai et al., 2007a; Tsujita et al., 2010; Chan et al., 2012) (Fig. 6F). Coimmunoprecipitation assays showed that the F-actin binding deficient Coro1A mutant (Coro1A^R29D^) was still able to coimmunoprecipitate myc-TRIMΔRING (Fig. 6G-H), suggesting that the Coro1A:TRIM67 interaction does not require Coro1A to be bound to F-actin. Coro1A binds the Arp2/3 complex in a phosphorylation dependent manner (Cai et al., 2005; Föger et al., 2006) (Fig. 6F). When Coro1A Serine 2 (Ser2) site is not phosphorylated, Coro1A binds to Arp2/3 complex, whereas phosphorylation of Ser2 via protein kinase C reduces the Coro1A interaction with the Arp2/3 complex. Both the phospho-mimetic (Coro1A^S2A^) and the non-phosphorylatable (Coro1A^S2D^) mutant of Coro1A were able to coimmunoprecipitate with and thus interact with TRIM67 (Fig. 6G-H). Taken together, the interaction between Coro1A and TRIM67 is independent of Coro1A binding to F-actin or phosphorylation of Ser2, which mediates Coro1A binding to the Arp2/3 complex.

Since we found no evidence that the Coro1A β-propeller was required for the interaction with TRIM67, we returned to the possibility that the coiled coil domain was responsible. We swapped either the coiled coil domain or the unique region between Coro1A and Coro1B (Fig. 6I), so that each construct maintains a trimerization domain. Coro1A bound TRIM67ΔRING strongly, but only a weak interaction was detected between TRIM67ΔRING and Coro1B. However, swapping the coiled coil domain between Coro1A and Coro1B reversed binding. Coro1A-1BCC coimmunoprecipitated a low amount of TRIM67ΔRING that was similar to Coro1B, whereas Coro1B-1ACC coimmunoprecipitated a high amount of TRIM67ΔRING that was similar to Coro1A. In addition, swapping the unique region did not affect Coro1A or Coro1B binding ability to TRIM67ΔRING (Fig. 6J-K). These data demonstrate that the coiled coil domain of Coro1A is critical for binding to TRIM67.

### The interaction between Coro1A and TRIM67 is required for netrin-dependent neuronal morphogenesis

We next explored the functional impact of losing the Coro1A:TRIM67 interaction during neuronal morphogenesis. To do so, we performed rescue experiments to evaluate neuronal responses to netrin in *Coro1a*^-/-^ neurons by reintroducing GFP-tagged wildtype Coro1A, Coro1B, or the Coro1A/1B coiled coil chimeras (Fig. 7A). Only wildtype Coro1A-GFP expression rescued the netrin-induced increase of growth cone area (Fig. 7B-C). Of note, *Coro1a*^-/-^ neurons had aberrantly large growth cones (Fig. 4). Introduction of Coro1A-GFP restored the *Coro1a*^-/-^ growth cone to a smaller size. Introduction of Coro1A-1BCC-GFP failed to rescue this phenotype or the netrin response. This suggests that the Coro1A coiled coil domain mediated Coro1A:TRIM67 interaction was required for proper growth cone morphology and responses to netrin. Intriguingly, both wildtype Coro1B-GFP and Coro1B-1ACC-GFP reduced growth size, indicating the Coro1B N-terminal region could also tune the growth cone morphology that is independent of TRIM67, but neither rescued response to netrin.

**Figure 7.**
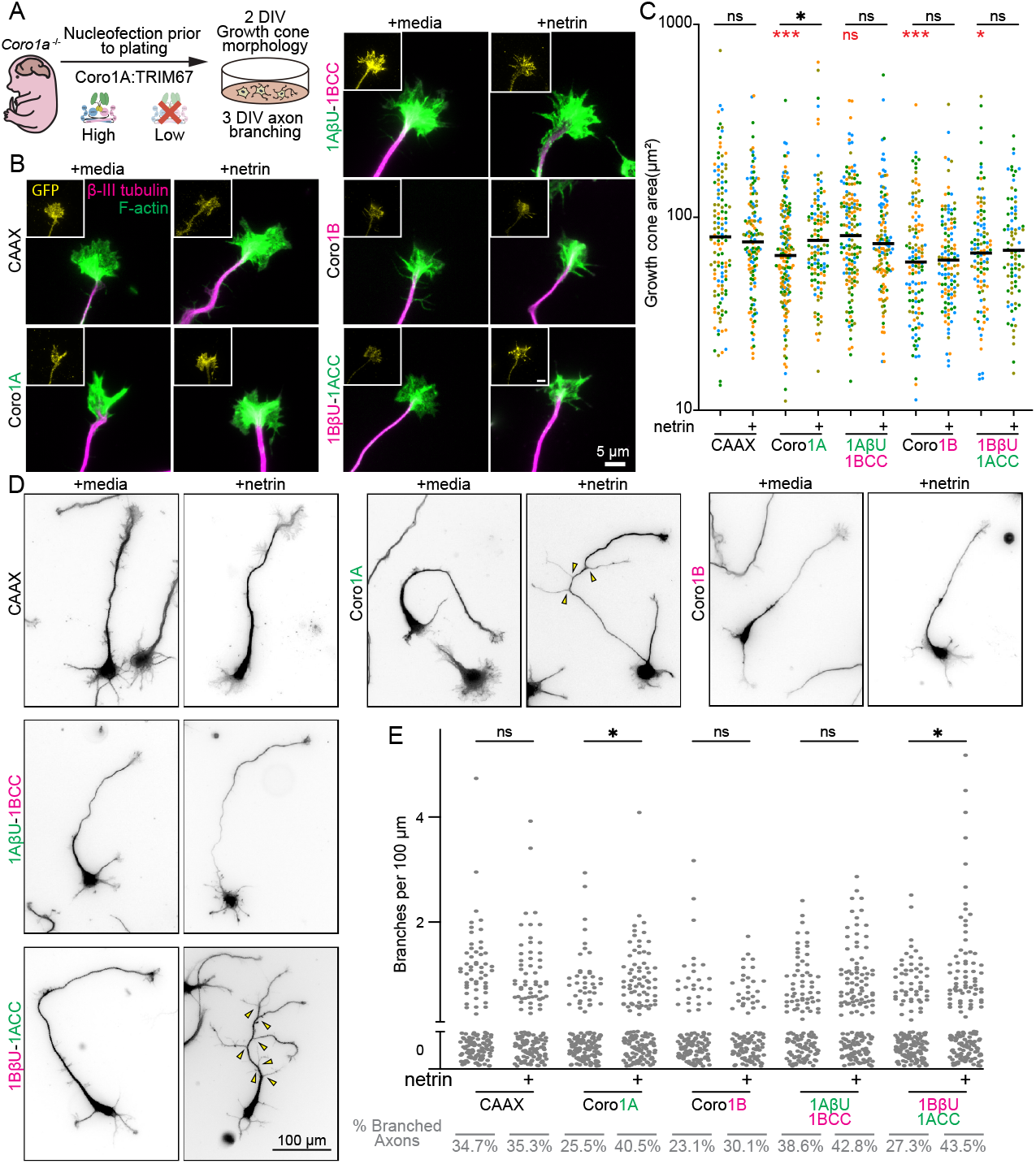
The Coro1A:TRIM67 interaction is required for netrin-dependent neuronal morphogenesis. (**A**) Schematic of experimental setup for rescue experiments. Rescue experiments were performed by expressing coronin wildtype or chimera constructs that either have strong or low binding to TRIM67. (**B**) Representative images of 2 DIV axonal growth cones from E15.5 *Coro1a*^-/-^ littermates expressing the indicated GFP-tagged constructs. Neurons were treated with media or netrin (600 ng/mL) for 40 min, and stained for GFP, β-III-tubulin and filamentous actin (phalloidin). (**C**) Quantification of growth cone size. n(cells)= 125 CAAX media, 133 CAAX netrin, 159 Coro1A media, 118 Coro1A netrin, 158 Coro1A-1BCC media, 127 Coro1A-1BCC netrin, 121 Coro1B media, 120 Coro1B netrin, 107 Coro1B-1ACC media, 84 Coro1B-1ACC netrin from three or four biological replicates. Paired two-tailed t-test was performed to compare growth size ± netrin (pairwise comparisons labeled in black), while one-way ANOVA (Bonferroni’s multiple comparisons tests) was performed to compare baseline growth cone size to the CAAX baseline condition (pairwise comparisons labeled in red). *, p <0.05; ***, p <0.001; ns, not statistically different. (**D**) Representative images of 3 DIV neurons from E15.5 *Coro1a*^-/-^ littermates expressing the indicated GFP-tagged constructs. Neurons were treated with media or netrin (250 ng/mL) for 24 hours, and stained for GFP, β-III-tubulin, and filamentous actin (phalloidin). (**E**) Quantification of axonal branching density per 100 µm axon length. n(cells)= 147 CAAX media, 152 CAAX netrin, 149 Coro1A media, 160 Coro1A netrin, 133 Coro1A-1BCC media, 154 Coro1A-1BCC netrin, 120 Coro1B media, 111 Coro1B netrin, 163 Coro1B-1ACC media, 155 Coro1B-1ACC netrin from four or five biological replicates. Kruskal-Wallis nonparametric ANOVA test (Dunn’s multiple comparison). *, p <0.05; ns, not statistically different.

For axon branching, both wildtype Coro1A-GFP and Coro1B-1ACC-GFP, which had high TRIM67 binding, rescued axon branching in response to netrin (Fig. 7D-E). Conversely, the chimeric proteins that had weak interaction with TRIM67, Coro1B-GFP and Coro1A-1BCC-GFP, failed to rescue axon branching (Fig. 7D-E). This suggests that the Coro1A coiled coil interaction with TRIM67 is essential for netrin-dependent axon branching. We conclude that netrin-dependent neuronal morphogenesis requires TRIM67 and Coro1A, and their interaction is crucial for this process.

## Discussion

Here we show that the actin binding protein Coro1A is a key regulator for neuronal morphogenesis. We found that Coro1A localized to the neuronal growth cone and was enriched at the base of filopodia, which are bundled actin protrusions critical for axon guidance and branching. We demonstrated that Coro1A was required for netrin-dependent axon guidance, growth cone responses, and axon branching in vitro. Consistent with a role in response to netrin, loss of Coro1A caused thinning of the corpus callosum in vivo. Furthermore, loss of Coro1A resulted in aberrantly large growth cones. Our previous work demonstrated that Coro1A interacts with TRIM67 (Menon et al., 2021), a brain-enriched E3 ubiquitin ligase that regulates neuronal responses to netrin. Interestingly, loss of Coro1A demonstrated similar phenotypes to loss of TRIM67, suggesting that Coro1A and TRIM67 are in the same netrin response pathway. We found that the coiled coil domain of Coro1A is essential for the Coro1A:TRIM67 interaction and showed that Coro1A:TRIM67 interaction was critical for netrin-dependent responses, indicating that Coro1A and TRIM67 collaborate in netrin-dependent neuronal morphogenesis.

### Coro1A is required for netrin-dependent axon guidance and branching

Our data support the hypothesis that Coro1A is critical for neuronal development. *CORO1A* is located in the 16p11.2 region of the human chromosome, a genomic locus where copy number variations are associated with neurodevelopmental and psychiatric disorders (Weiss et al., 2008; McCarthy et al., 2009; Niarchou et al., 2019). Many genes within this region are required for normal nervous system development, including *MAPK3* (Pucilowska et al., 2015), *TAOK2* (Yadav et al., 2017), and *KCTD13* (Escamilla et al., 2017; Kizner et al., 2020), with our data implicating Coro1A as well. Others have shown that Coro1A is required for proper axon guidance in vivo (Blaker-Lee et al., 2012). Knocking down *coro1a* in zebrafish resulted in disorganized axon tracts in both the forebrain and hindbrain. Of note, zebrafish possess a single copy of *coro1a*. Similarly, we find that Coro1A is required for axon guidance under a netrin gradient. One key question is how Coro1A regulates axon guidance. Axon guidance requires precise spatiotemporal remodeling of the actin cytoskeleton in the growth cone. Coronins are F-actin regulators in non-neuronal cells. We showed that Coro1A is similarly enriched in neuronal growth cones, where it regulates the actin cytoskeleton. Our results show without netrin, that *Coro1a*^-/-^ neurons exhibit enlarged growth cones. This phenotype may result from increased Arp2/3 activity or reduced actin debranching, as Coro1A has been shown to inhibit the activation of Arp2/3 complex and promote actin debranching. Coronin also promotes cofilin-mediated actin disassembly in non-neuronal cells. Therefore, the loss of Coro1A may slow cofilin-mediated disassembly, contributing to enlarged growth cones. Disentangling the inhibitory function of Coro1A on Arp2/3 and its synergistic role in cofilin is challenging as both alter actin turnover. Interestingly, although Coro1B expression in *Coro1a*^-/-^ neurons was unable to rescue growth cone response to netrin, it did rescue the enlarged growth cone phenotype caused be loss of Coro1A. This suggests potential overlapping functions for Coro1A and Coro1B in maintaining the F-actin network within the growth cones while highlighting the specificity of Coro1A in TRIM67 and netrin-dependent signaling pathways.

We discovered that Coro1A is required for netrin-dependent axon branching. One proposed model for axon branch formation begins with the formation of Arp2/3-based F-actin patches along the axon (Ketschek and Gallo, 2010; Spillane et al., 2011, 2012). These patches serve as sites of de novo filopodia emergence that eventually stabilize to form branches. Coro1A may contribute to the formation or stabilization of actin patches by directly interacting with Arp2/3 or F-actin. Alternatively, Coro1A localizing to the axonal filopodia suggests it may be involved in regulating the filopodia stability and lifetime, potentially promoting their transition into stable branches. Alternatively, or in addition, Coro1A may play a role in the convergence of branched actin into linear actin and thus the emergence of filopodia, which is critical step driven by actin binding proteins like VASP and formins. Our recent work shows that TRIM67 and TRIM9 interact with the actin polymerase VASP and antagonistically regulate its non-degradative ubiquitination (Boyer et al., 2020), thereby modulating VASP function and actin dynamics that are important for netrin responses (McCormick et al., 2024b). We were surprised not to find evidence of ubiquitin-mediated regulation of Coro1A here. This indicates that there are likely multiple mechanisms by which TRIM proteins can regulate protein functions during netrin responses beyond ubiquitylation.

### Coro1A role in synaptogenesis

Proteomic studies from us and others have detected Coro1A in the PSD (Gay et al., 2024; Mccormick et al., 2024; McCormick et al., 2024a). Although our work focused on Coro1A in early neuronal development prior to synaptogenesis, we observed increased levels of Coro1A in later stages of neuron development in vivo and verified that Coro1A was enriched in the PSD fraction (Fig. S1C). These findings are consistent with Coro1A regulating synaptogenesis and plasticity. Coro1A is enriched in excitatory synapses and regulates the excitatory/inhibitory synapse ratio (Jayachandran et al., 2014), yet whether Coro1A plays a role in dendritic spine morphology, function, or actin architecture is unknown. Like the leading edge of migrating cells and neuronal growth cones, branched actin networks regulated by both Arp2/3 complex and cofilin are critical for dendritic spine maturation and plasticity (Hotulainen and Hoogenraad, 2010; Borovac et al., 2018; Spence and Soderling, 2015; Konietzny et al., 2017). Thus, we expect Coro1A may be poised to tune the branched actin network within the dendritic spine head that could be key to spine morphology, maturation, and function. Recent studies from our lab and others have shown that netrin promotes the formation of dendritic filipodia and spines, neuronal activity, and synaptic plasticity (Goldman et al., 2013; Glasgow et al., 2018; Wong et al., 2019; McCormick et al., 2024a). In addition, our previous work demonstrates that TRIM67 is also enriched in the PSD fraction (Mccormick et al., 2024). Whether a common collaborative mechanism of Coro1A and TRIM67 is involved for netrin responses during later stages of neuronal development such as synaptogenesis and synapse maturation is intriguing.

### Beyond trimerization of the coronin coiled coil domain

The coiled coil domain of coronins mediates coronin oligomerization. Some studies have shown that the coiled coil domain of yeast coronin (Crn1) mediates direct binding to and inhibition of the Arp2/3 complex (Humphries et al., 2002; Rodal et al., 2005; Sokolova et al., 2017). However, other studies suggest that Crn1 binding to the Arp2/3 complex is mediated by the unique region (Liu et al., 2011; Koundinya et al., 2025). Interestingly, our analysis revealed that the coiled coil domain of Coro1A is critical for TRIM67 binding. In addition, the Coro1A:TRIM67 is essential for netrin-dependent neuronal shape changes, underlying a possible additional function of the coiled coil domain beyond coronin oligomerization. One possibility is that Coro1A could serve as a scaffold to target TRIM67 to specific regions or structures within the cell. However, whether Coro1A oligomerization and TRIM67 binding are mutually exclusive remains unknown. Coro1A may co-oligomerize with TRIM67, or alternatively TRIM67 may interact with a binding interface existing on the exterior of the coiled coil domain of Coro1A. Future structural studies will be necessary to understand this more fully. Supporting the idea that the coiled coil domain is not merely a trimerization domain, a recent study showed that triple knockdown of Coro1A, Coro1B, and Coro1C in COS7 cells resulted in extended actin on endosome buds, decreased endosome fission rate, and ER tubule contact with endosomes (Striepen and Voeltz, 2022). Notably, expression of the coiled coil domain of Coro1C alone was sufficient to rescue all these phenotypes, highlighting a potentially essential role of a Coro1C coiled coil domain beyond oligomerization.

### A role for Coro1A in filopodia

Type I coronins are well-recognized branched actin regulators. Interestingly, coronins are also capable of binding linear F-actin in vitro (Jansen et al., 2015), and a recent overexpression study in U2OS cells demonstrated Coro1B localizes to the filopodia shaft (Jacquemet et al., 2019). These findings raise the question whether endogenous coronin localizes to filopodia. Our discoveries provide new insights, as we found endogenous and overexpressed Coro1A localizes to the base of neuronal filopodia, suggesting that Coro1A is not only a branched actin regulator, but also plays a role in filopodia dynamics or structure in neurons. One potential function could be bundling actin filaments, as biochemistry studies with yeast Crn1 have demonstrated an actin crosslinking effect (Goode et al., 1999; Han et al., 2023). Alternatively, Coro1A binding to filaments might alter ultrastructure or induce twisting. Another hypothesis is that Coro1A may prevent or recruit specific actin binding proteins or non-actin binding proteins to the base of the filopodia, such as cofilin and TRIM67, respectively. Previous in vitro work suggested that Coro1B binds to filaments and subsequently recruits cofilin for severing (Jansen et al., 2015). However, a recent study showed the loss of coronins actually resulted in an enrichment of cofilin in the lamellipodia (King et al., 2022), which in turn stabilized actin filaments due to cofilin saturation and reduced severing efficiency. Future examination of neuronal filopodia is important to characterize the coronin/cofilin regulatory pathways. Finally, the filopodia base localization pattern is reminiscent of cofilactin recently described in growth cones (Hylton et al., 2022; Minamide et al., 2023), which are cofilin-coated actin at the base of filopodia. The cofilactin filaments alter the packing and ultrastructure of actin bundles, actin filaments are hyper-twisted within these regions and are excluded from fascin-binding. Given the interplay between coronins and cofilin, an intriguing question would be whether Coro1A plays a role in mediating cofilactin including length and dynamics.

## Material and methods

### Animals

All mouse lines were on a mixed C57BL/6J background and bred at the University of North Carolina with approval from the Institutional Animal Care and Use Committee. Generation of *Coro1a*^-/-^ and *Trim67*^-/-^ mice were described (Föger et al., 2006; Boyer et al., 2018). Timed pregnant females were obtained by placing male and female mice together overnight; the following day was designated as embryonic day (E) 0.5 if the female had a vaginal plug. The sexes of postnatal day (P) 4 mice for corpus callosum assay were determined by checking for the presence of a pigmented spot on the scrotum prior to perfusions (Wolterink-Donselaar et al., 2009) and genotyping.

### Plasmids, antibodies, and reagents

Details of plasmids are included in Supplemental Table 2. Plasmids of myc-TRIM67 and myc-TRIM67ΔRING were described (Boyer et al., 2018). GFP tagged Coro1AΔCC, Coro1AΔUC, Coro1AΔU, Coro1A^R29D^, Coro1A^S2D^, Coro1A^S2A^ constructs were generated by NEB Q5 site-directed mutagenesis or QuikChange mutagenesis of Coro1A-GFP plasmid. Coro1A gene fragment with all the unique residues mutated to Coro1B or Coro1C was made by IDT and then cloned into GFP vector to make the mutated Coro1A surface residue construct. Chimeras of Coro1A and Coro1B were created by NEB HiFi DNA assembly. Myc-TRIM67ΔRING domain deletion constructs were generated by NEB Q5 site-directed mutagenesis of myc-TRIM67 domain deletion constructs described previously (Boyer et al., 2020). Myc-TRIM67 fragments 159-279, 280-490, and 489-761 were generated by NEB Q5 site-directed mutagenesis or HiFi DNA assembly of myc-TRIM67ΔRING. The following plasmids were acquired: Coro1A-GFP, Coro1B-GFP and Coro1C-GFP (James Bear, University of North Carolina at Chapel Hill), pRK5-HA-Ubiquitin-WT and pRK5-HA-Ubiquitin-K0 (Lim et al., 2005) were gifts from Ted Dawson (Johns Hopkins University, Baltimore, MD; Addgene plasmid 17608, RRID:Addgene_- 17608; Addgene plasmid 17603, RRID:Addgene_- 17603).

Antibodies used were: mouse monoclonal against c-Myc (9E10) [Hybridoma serum purified in-house; 1:10000 for western blotting (WB)], rabbit polyclonal against GFP (A11122, Invitrogen; 1:3000 WB), rabbit polyclonal against TRIM67 [generated in house (Boyer et al., 2018); 1:2000 WB], chicken polyclonal against GFP [GFP-1010, Aves Labs; 1:5000 WB and 1:500 immunofluorescence staining (IF)], rabbit monoclonal against Coro1A (ab203698, Abcam; 1:1000 WB and 1:100 IF), GAPDH (sc-166545, SCBT; 1:1000 WB), mouse monoclonal against Coro1A (sc-100925, SCBT; 1:1000 WB), rabbit polyclonal against β-III-tubulin (Tuj1, 802001, BioLegend; WB 1:5000), mouse monoclonal against Fascin 1 (sc-21743, SCBT; 1:500 IF), rat monoclonal against L1CAM (Sigma-Aldrich, MAB5272; 1:500 IF), mouse monoclonal β-III Tubulin (801202, BioLegend; IF 1:4000), mouse monoclonal against HA (12CA5) (Hybridoma serum, Patrick Brennwald, University of North Carolina; 1:100 WB), mouse monoclonal against GFP (75-131, University of California, Davis, Neuromab; immunoprecipitation 1:1500), mouse monoclonal PSD-95 (75-028, University of California, Davis, Neuromab; 1:500,000 WB).

Adenovirus expressing GFP-actin was a generous gift from James Bamburg, Colorado State University. Doxycycline-inducible adenoviral constructs were generated for Coro1A-GFP and HA-ubiquitin using the Adeno-X system 3 (631180, Takara Bio).

Recombinant netrin was concentrated as previously described (Serafini et al., 1996). Briefly, HEK293 cells stably expressing chicken netrin were cultured in growth media [DMEM (11965092, Thermo Fischer Scientific) supplemented with 10% FBS (MP1300500, VWR), 250 ng/mL Geneticin (10131035, Thermo Fischer Scientific), 200 µg/mL Hygromycin (30-240-CR, Corning), 1% Penicillin-Streptomycin (15140122, Thermo Fischer Scientific)] to 80-90% confluency. Cells were washed with warm PBS containing calcium and magnesium, replaced with Opti-MEM (11058021, Thermo Fischer Scientific) and incubated for 16-24 hrs at 5% CO2/37°C. The next day, Opti-MEM containing secreted netrin was concentrated using Amicon filters tubes with a 30 kDa cut-off (UFC903008, Millipore). Netrin concentration was determined using SDS-PAGE and Coomassie staining.

### Cortical neuron culture, transfections, and transductions

Cortical neuron cultures were prepared from E15.5 mice. Cortices were dissected in HEPES/HBSS buffer, digested with 0.25% trypsin (25-054-CI, Corning) for 20 min at 37 °C, washed three times with trypsin quenching media [TQM: neurobasal media (21103049, Thermo Fischer Scientific) supplemented with 2 mM Glutamax (35050061, Thermo Fischer Scientific) and 5% fetal bovine serum (MP1300500, VWR)]. The washed cortices were gently triturated with a P1000 pipette tip, counted by hemocytometer, and plated in trypsin quenching media onto coverslips or dishes coated with 1 mg/mL poly-d-lysine (P7886, Sigma-Aldrich). If embryos from *Coro1a*^+/-^ pregnant mice were used, the dissected brains were stored at 4°C in HibernateE (A1247601, Thermo Fischer Scientific) supplemented with 2mM Glutamax and 2% B27 (17504044, Thermo Fischer Scientific) until genotypes were confirmed. One to three hours after plating, the media was changed into serum free media [SFM: neurobasal media supplemented with 2% B27 and 2 mM Glutamax]. Neuron cultures were maintained at 5% CO2/37°C. For growth cone morphology assays, at 2 DIV either serum free media or 600 ng/mL of netrin was added to the culture 40 min prior to fixing cells. For axon branching assays, at 2 DIV either serum free media or 250 ng/mL of netrin was added to culture. Neurons were then fixed 24h after treatment.

For rescue experiments, Amaxa Nucleofector was used to transfect neurons prior to plating. Briefly, dissociated neurons were pelleted at 100 x g for 7 min, resuspended in Nucleofector solution (VPG-1001, Lonza) with 1.5-2 µg of the plasmid of interest, and electroporated with a nucleofector according to the manufacturer protocol. For neuron immunoprecipitation assays and examining Coro1A dynamics for live-cell imaging, concentrated adenovirus was added to each dish one to three hours after plating. If the adenovirus expressed a doxycycline-inducible construct, 1 µg/mL doxycycline was added 48 h prior to the experiment endpoint.

### HEK293 cell culture and transfection

Wildtype HEK293 cells (female) were obtained from Simon Rothenfußer (Klinikum der Universität München, München, Germany). *TRIM67*^-/-^ HEK293 cells were generated as previously described (Boyer et al., 2020). HEK293 cells were grown at 5% CO2/37°C in DMEM with glutamine (11965092, Thermo Fischer Scientific), supplemented with 10% FBS (MP1300500, VWR). HEK293 cells were transfected using Polyplus jetPRIME reagent (55-132, Genessee) according to the manufacturer’s protocol. Briefly, 1.5 -2 µg of each plasmid was mixed with 200 µL jetPRIME buffer and 8 µL jetPRIME. The solution was vortexed for 10s, spun down, and incubated for 10 min at room temperature. The solution was then gently added dropwise to HEK293 cells cultured in 60 mm dishes at 70% confluency.

### Western blotting

All samples were resolved by SDS-PAGE and transferred onto 0.45 µm nitrocellulose membrane for immunoblotting. Membranes were blocked in 5% milk in Tris-buffered saline (TBS) for 1h at RT and then probed with primary antibodies diluted in 2% milk in TBS with 0.1% Tween-20 (TBST) overnight at 4°C. The next day, membranes were washed three times with TBST, incubated with secondary antibodies diluted in 2% milk in TBST, washed again three times with TBST, and imaged using an Odyssey Imager (LI-COR Biosciences).

### Coimmunoprecipitation assays

For coimmunoprecipitation assays in HEK293 cells, 16-18 hours post-transfection cells were lysed in immunoprecipitation (IP) buffer (10% glycerol, 50 mM Tris at pH7.5, 200 mM NaCl, 2 mM MgCl2, 1% NP-40, supplemented with 15 mM sodium pyrophosphate, 50 mM sodium fluoride, 40 mM β-glycophosphate, 1 mM sodium vanadate, 100 mM phenylmethylsulfonylfluoride, 1 µg/mL leupeptin, and 5 µg/mL aprotinin) for 10 minutes on ice. Cells were scraped off the dish, transferred into tubes and centrifuged at 18,400 xg for 15 min at 4°C. Concentration of the solubilized protein was determined by Bradford assay and then normalized before being subjected to immunoprecipitation using GFP-TRAP (gtma, proteintech) or myc-TRAP (ytma, proteintech) beads. For each condition, a total of 500-1000ug of lysate at a concentration of 1 mg/mL were rotated with the 10 µL beads for 1h at 4°C. Beads were then collected, washed three times with IP buffer, and then boiled with 2X sample buffer (65.8 mM Tris-HCl, pH 6.8, 2.1% SDS, 26.3% glycerol, 0.01% bromophenol blue) containing β-mercaptoethanol.

For coimmunoprecipitation assays in cultured neurons, 10 million neurons were plated and infected with adenovirus expressing Coro1A-GFP 1h after plating. At 3 DIV, neurons were lysed and subjected to the same GFP-TRAP protocol as described for HEK293 cells, except that 10 µM Latrunculin B (L5288, Sigma-Aldrich,) was supplemented in the IP buffer.

### Analysis of protein lysate

Cortices from *Coro1a*^+/+^ and *Coro1a*^-/-^ littermates were dissected out from developmental time points and flash frozen. Frozen cortices were lysed in ice-cold IP buffer for 30 min, homogenized with a dounce homogenizer, and centrifuged at 18,400 xg for 15 min. Following protein concentration measurements via Bradford assays, supernatants were boiled with 2X sample buffer containing β-mercaptoethanol. For measuring Coro1A protein level changes in cultured neurons, neurons were lysed in RIPA buffer (50 mM Tris, pH 8, 150 mM NaCl, 0.5% deoxycholate, 0.1% SDS, 1% NP-40 supplemented with 15 mM sodium pyrophosphate, 50 mM sodium fluoride, 40 mM β-glycophosphate, 1 mM sodium vanadate, 100 mM phenylmethylsulfonylfluoride, 1 µg/mL leupeptin, and 5 µg/mL aprotinin) and centrifuged at 18,400 xg for 15 min. Concentration of supernatant was measured, and the supernatant were then boiled with 2X sample buffer containing β-mercaptoethanol.

### Postsynaptic density preparation

The forebrain (cortex and hippocampus) of three-week-old *Coro1a*^+/+^ and *Coro1a*^-/-^ littermates were dissected out in ice-cold dissection media, and flash frozen in liquid nitrogen. Frozen forebrains were homogenized using 15 strokes from a homogenizer in ice cold homogenization buffer (10 mM HEPES pH 7.4, 320 mM sucrose, 1 mM EDTA, 5 mM sodium pyrophosphate, 1 mM sodium vanadate, 150 µg/ml phenylmethylsulfonyl fluoride, 2 µg/ml leupeptin, 2 µg/ml aprotinin). The brain homogenate (HOM) was next centrifuged at 800 × g for 10 min to remove nuclear fraction (pellet 1). The supernatant (S1) was then centrifuged at 16,000 × g. The pellet (pellet 2) was resuspended in homogenization buffer, layered onto a discontinuous sucrose gradient (0.8M, 1M or 1.2M sucrose plus protease and phosphatase inhibitors mentioned above), and then subjected to ultracentrifugation at 82.5 k × g for 90 min. The material at the interface between 1.2 and 1 M sucrose (synaptosomes) was collected. Synaptosomes were diluted with 10mM HEPES buffer (plus protease and phosphatase inhibitors mentioned above) to reduce sucrose concentration back to 320 mM, and pelleted at 100 k × g for 30 min. The pellet was resuspended in 50mM HEPES buffer (plus protease and phosphatase inhibitors mentioned above) and then mixed with an equal part of 1% Triton-X-100. The mixture was incubated for 15 min rotating, centrifuged at 32 k × g for 20 min. The pellet (PSD fraction) was resuspended in 50mM HEPES buffer (plus protease and phosphatase inhibitors). Sample concentrations were determined by Bradford assays, then boiled with 2X sample buffer containing β-mercaptoethanol.

### Cultured neurons fixation and immunostaining

For fixed-cell imaging experiments, neurons were cultured on 1 mg/mL poly-d-lysine-coated #1.5 German glass coverslips (72290-04, ElectronMicroscopy Sciences). Prior to coating, coverslips were cleaned using Harrick Plasma Cleaner (PDC-32G). For growth cone morphology and axon branching assays, neurons were fixed with warm 4% paraformaldehyde in PHEM buffer (60 mM PIPES, 25 mM HEPES, 10 mM EGTA, 2 mM MgSO4, and 0.12 M sucrose) for 20 min at RT, rinsed three times with 1X PBS, and then permeabilized with 0.1% Triton X-100 for 10 min. To examine Coro1A sub-filopodia localization, an alternative fixation method used for visualizing cofilactin rods was performed (Hylton et al., 2022; Minamide et al., 2023). These neurons were simultaneously fixed and permeabilized with icecold methanol at -20°C for 20 min followed by three 1X PBS rinses.

For immunostaining, permeabilized cells were blocked in 10% donkey serum or 3% bovine serum albumin (BSA) for 30 min at RT, stained with primary antibodies diluted in 1% donkey serum or BSA for 1h at RT, rinsed three times with 1X PBS, stained with secondary antibodies diluted in 1% donkey serum or BSA for 1h at RT, washed three times with 1X PBS, and mounted with Tris/glycerol/n-propyl-gallate–based mounting media. A 30 min phalloidin stain (diluted in 1X PBS) was performed after secondary antibody washes.

### Growth cone turning assays

Microfluidic devices were prepared as previously described (Taylor et al., 2015; Boyer et al., 2020; Mutalik et al., 2025). Briefly, Polydimethylsiloxane (PDMS) (Sylgard 184, Dow Corning) and curing agent were mixed in a 10:1 (by weight) ratio. 10g of the mixture was poured onto each silicon wafer, degassed in a desiccator for 2-3h, and then cured at 60°C for at least overnight. A day before plating, the devices were cut out, and the edge of each fluid chambers were punched. Each individual devices were cleaned with 3M tape, soaked in 70% ethanol, and air dried for 1h before attaching to PDL-coated coverslip. SFM supplemented with anti-anti were added to all chambers, then devices were stored in a 5% CO2/37°C incubator for overnight. For the coverslips (48404-467, VWR), a day prior to assembling devices coverslips were soaked in 100% ethanol and bath sonicated for 30 m. After the sonication, coverslips were dried for 1h, transferred into PDL solution (354210, Corning), and then incubated overnight at 37°C. The next day, coverslips were rinsed with MiliQ water twice and dried for at least 1h. For plating, 10 µL of dissociated *Coro1a*^+/+^ or *Coro1a*^-/-^ E15.5 neurons (18 million/mL) were added to the cell compartment. Once the axons reached the axon viewing chamber (48h-72h), either a control dextran gradient or dextran+netrin (600 ng/mL) gradient was established. Gradients were established by adding dextran/netrin solutions to one side of the source chamber and removing 50 µL media from the upper sink chamber. Immediately after the gradient was established, the devices were placed on the microscope for imaging. DIC or phase contrast (axons) images were taken every 5 min, and epifluorescence (dextran) images to confirm the gradient were acquired every 1h.

### Neuroanatomic imaging

P4 *Coro1a*^+/+^ and *Coro1a*^-/-^ littermates were anesthetized with an intraperitoneal injection of 1.2% avertin and transcardially perfused with 4% PFA in PBS. Brains were removed and post-fixed in 4% PFA for 48h at 4°C. Brains were washed three times with 1X PBS, incubated in 30% sucrose in PBS at 4°C for 2 days, and embedded in O.C.T. Compound (4585, Fisher). 20 µm coronal sections were collected every 100 µm anterior to posterior. Sections were permeabilized in 0.3% Triton-X in PBS for 30 min at RT, then blocked in 5% normal goat serum in PBS for 2h at RT. Primary antibody against L1CAM diluted in 5% normal goat serum was then applied to the sections overnight at 4°C. After three rinses with PBS, sections were stained with Alexa Fluor PLUS 647 Goat anti-Rat IgG diluted in 5% normal goat serum for 30 min at RT. Then they were washed again three times with PBS, and stained with DAPI for 15 min. Finally, the sections were mounted with Fluoromount G (004958-02, Invitrogen) and imaged with a 10X/0.4-NA objective on an Olympus VS200 Slide Scannerslide scanner.

### Ubiquitination assays

The ubiquitination assays in HEK293 cells were performed as described (Urbina et al., 2021; Mutalik et al., 2025). Briefly, HEK293 cells were transfected with Coro1A-GFP and HA-ubiquitin or HA-ubiquitin^K0^ using Polyplus jetPRIME reagent. 16-18 hours posttransfection cells were treated with 10 µM MG132 for 1h and lysed in ubiquitination immunoprecipitation buffer (50 mM Tris at pH7.5, 150 mM NaCl, 1 mM EDTA, 0.5% Triton X, 5 mM N-ethylmaleimide, 50 µM PR619, supplemented with 15 mM sodium pyrophosphate, 50 mM sodium fluoride, 40 mM β-glycophosphate, 1 mM sodium vanadate, 100 mM phenylmethylsulfonylfluoride, 1 µg/mL leupeptin, and 5 µg/mL aprotinin). Lysates were collected and centrifuged at 18,400 xg for 10 min at 4°C. Lysates were diluted to 1 mg/mL. 1000 µg of protein were incubated with 15 µL GFP-TRAP beads for 2h at 4°C on a rotator. Beads were washed with wash buffer (10 mM Tris at pH7.5, 150 mM NaCl, 0.5 mM EDTA, 10 mM N-ethylmaleimide, 50 µM PR619, supplemented with 15 mM sodium pyrophosphate, 50 mM NaF, 40 mM β-glycophosphate, 100 mM phenylmethylsulfonylfluoride, 1 mM sodium vanadate, 1 µg/mL leupeptin, 5 µg/mL aprotinin), followed by three washes with stringent wash buffer (8M urea, 1% SDS, in 1X PBS), and a SDS wash buffer (1% SDS in 1X PBS). Beads were boiled with 2X sample buffer containing β-mercaptoethanol.

Ubiquitination assays in cultured neurons were performed similarly as described previously (Menon et al., 2015; Boyer et al., 2020). Briefly, 10 million *Trim67* ^+/+^ and *Trim67*^-/-^ neurons were plated and transduced with adenovirus expressing Coro1A-GFP and HA-ubiquitin 1h after plating. At 3 DIV, neurons were treated with 10 µM MG132 for 2h and lysed in 200 µL neuron ubiquitination buffer (20 mM Tris at pH7.5, 250 mM NaCl, 3 mM EDTA, 3 mM EGTA, 0.5% NP-40, 1% SDS, 2 mM dithiothreitol, 5 mM N-ethylmaleimide, 50 µM PR619, supplemented with 15 mM sodium pyrophosphate, 50 mM sodium fluoride, 40 mM β-glycophosphate, 1 mM sodium vanadate, 100 mM phenylmethylsulfonylfluoride, 1 µg/mL leupeptin, and 5 µg/mL aprotinin). For netrin treated groups, 600 ng/mL netrin was added 1h before lysing. Lysates were gently vortexed, boiled for 20 min, centrifuged at 18,400 xg for 10 min and diluted with neuron ubiquitination buffer without SDS so that final concentration of SDS was reduced to 0.1%. Protein concentrations were determined by Bradford assay, and each sample was diluted to the same concentration. Mouse monoclonal antibody against GFP was added to each sample and incubated overnight at 4°C on a rotator. The next day, protein A beads were added and incubated for 2h to capture antibody-bound substrates. Beads were next washed with neuron ubiquitination buffer twice and then boiled with 2X sample buffer containing β-mercaptoethanol.

### Coro1A live-cell imaging and FRAP imaging

For Coro1A dynamic assays, neurons expressing Coro1A-GFP were imaged at 2 DIV. TIRF images were acquired every 2s for 2 min with 200 ms exposure with 491 laser with 30% power.

For FRAP assays, neurons expressing GFP-actin were imaged at 2 DIV. Prebleach images were acquired at 130ms intervals for 5 frames with 50ms exposure with 491 laser with 30% power. Using 405nm laser, neurons were bleached for 50 ms to acquire FRAP ROI with camera, then immediately bleached for 200ms with 405nm with 100% power. Neurons were then imaged for 600 frames at 130 ms intervals with 50 ms exposures of 491 nm laser with 30

### Microscope descriptions

Endogenous and exogenous Coro1A localization experiments, growth cone morphology assays, axon turning assays and FRAP experiments were performed on an Olympus IX83-ZDC2 equipped with a cMOS camera (Orca-Fusion, Hamamatsu), Xenon light source, and the following objective lenses: a UPlanApo 100×/1.5-NA DIC TIRF objective (Olympus), a UAPON 100×/1.49-NA DIC TIRF objective (Olympus), a 20×/0.85-NA UPlanSApo DIC objective lens (Olympus), using Cellsens software (Olympus) or Metamorph acquisition software. Cells for live imaging were maintained in culture medium and at 37°C and 5% CO2 with a Tokai-Hit on-stage cell incubator. Axon branching and axon turning assays were conducted on Keyence BZ-X800 All-in-one Fluorescence Microscope using a Keyence Plan Apochromat 40×/0.95-NA objective and Keyence Plan Fluorite 10×/0.3-NA objective, respectively. Cells for live imaging were maintained in culture medium and at 37°C and 5% CO2 with a TokaiHit on-stage cell incubator. Coronal brain sections were acquired on an Olympus VS200 Slide Scannerslide scanner equipped with a 10X/0.4-NA objective.

### Western blot analysis

Western blots were quantified using Li-Cor Image Studio Lite. Background was subtracted for each quantified band. For coimmunoprecipitation assays, the coimmunoprecipitated band intensity was normalized to the immunoprecipitated band intensity, then the ratios were log-transformed followed by statistical analysis. To compare protein level changes across development, the Coro1A or TRIM67 measured intensity was normalized to the loading control (β-III-tubulin or GAPDH). To compare ubiquitin level across conditions, the sum of all HA-ubiquitin bands was divided by amount of Coro1A-GFP immunoprecipitated.

### Filopodia, growth cone and axon branching analysis

Coro1A filopodia distribution analysis was adapted from FiloMap (Popović et al., 2023; Jacquemet, 2023). Briefly, filopodia line scans were drawn manually from tip to base using the fascin channel in ImageJ. To normalize filopodia length variations, each line intensity profile was binned into 40 bins, with bin 1 corresponding to the tip and bin 40 to the base. The average intensity value within each bin assigned to that bin. The binned profiles were then pooled and averaged to create a heatmap for visualization.

For growth cone size measurements, images were first blinded in ImageJ and then imported into ImageTank for analysis using a custom script. Briefly, we first cropped the region of the image containing the growth cone. The intensity of the images was then scaled based on the minimum and maximum intensity. To identify the boundary of the growth cones a large median filter (radius 6) followed by sharpening (sigma 15, width 1.5) was applied to the image. A threshold value to segment the growth cones was set based on visual inspection and applied to images. The initial mask was modified but the morphological operation close (radius, 2) followed by selection of the largest continuous object. The area of the growth cone was defined as the area of the resulting mask. Two masks were considered. One filled the entire mask area and the other permitted holes within the mask. We defaulted to a filled mask but compared the areas determined by each approach and if the difference was greater than 5% we visually inspected the masks. This was because occasionally filopodia cross each other and the space between them should not be included in the area of the growth cone.

Axon branching analysis was performed in FIJI using the Neuroanatomy Simple Neurite Tracer plugin (Longair et al., 2011). All quantification of images was performed blind. Axon branches were defined as extensions off the primary axon that are greater than 20 µm. Axon branching density was defined as the number of branches per 100 µm of axon.

### Axon turning analysis

Axon turning was analyzed using a custom script in ImageTank described previously (Mutalik et al., 2025). Briefly, growth angle relative to a horizontal line drawn down the gradient was measured at the initial and final time frames. Axon turning angle was measured by calculating the difference between the growth angle of the start and end point. A positive angle indicated turning towards the gradient. To be included for analysis, neurons had to be healthy or trackable for at least 2hrs (tracking was terminated at 10hrs). In addition, axons that grew along other axons, retracted, or stalled (did not grow more than 5 µm) were excluded from analysis. Branches were included if they clearly separated from the primary axon and met the above criteria.

### FRAP analysis

To analyze FRAP experiments, all measurements were taken using a circle with a diameter of 4.03 µm. The intensities of the FRAP ROI, three separate areas of the growth cone were used to assess photofading (cell), and the background (bkgd) were measured. The following equation was used to correct for background fluorescence photofading, and loss of fluorescent material due to the bleach, and normalized to correct for differences in protein expression between neurons.

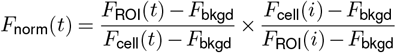

The normalized intensity of the FRAP ROI for each cell was fit to the following equation, where A is the fluorescence recovery plateau, τ is the recovery time constant, and t is time:

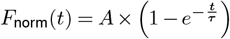

The plateau value was extracted from the curve, and fluorescent recovery halftime was derived from τ using:

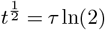

### AlphaFold

AlphaFold2 (Jumper et al., 2021, Evans et al., 2022) was used to predict molecular structures of TRIM67. TRIM67 structures containing different subsets of domains were predicted and analyzed. Both monomeric and dimeric structures were predicted. The predicted dimer structure containing amino acids 280-497 (BBox2-coiled coil-Cos) formed one long helix that coiled in an antiparallel conformation with the BBox2 domains stably positioned at each end of the long coiledcoil region.

### Statistics

All statistical analysis was completed in Prism10 (Graphpad). A minimum of three biological experiments were performed for data performed with statistics. For normally distributed data, t test was performed when two groups were compared, or one-way ANOVA with post-hoc correction (method stated in the figure legend) was performed for multiple comparisons. For nonnormal datasets, we used Kruskal–Wallis nonparametric ANOVA with Dunn’s correction for statistical analysis. Statistical differences were determined by an α= 0.05.

### Online Supplemental Material

Fig S1 shows RNAseq of type I coronins in developing rodent brain and Coro1A protein enrichment in the postsynaptic density fraction. Fig S2 validates specificity of Coro1A antibody. Fig S3 shows FRAP of GFP-actin in growth cones of *Coro1a*^+/+^ or *Coro1a*^-/-^ neurons. Fig S4 details mapping of the interaction between TRIM67 and Coro1A. Table 1 details mutations made on unique surface residues of Coro1A β-propeller. Table 2 provides details of plasmids used in study. Supplemental Video 1 and 2 demonstrate Coro1A-GFP dynamics in the developing neuron. Supplemental Video 3, 4, 5 demonstrate microfluidic based axon turning assays in a dextran (3), low netrin (4) or high netrin (5) gradient. Supplemental movies 6-9 demonstrate FRAP assays of GFP-actin in *Coro1a*^+/+^ or *Coro1a*^-/-^ growth cones.

## Acknowledgements

We thank Janee Cadlett-Jette and Natallia Riddick for assistance with mouse husbandry. We thank David Adalsteinsson for the generous help with the ImageTank software. We thank James Bamburg (Colorado State University) for providing the GFP-actin adenovirus for actin turnover experiments. We thank Laura McCormick and Charise White for the postsynaptic density preparation. This work was supported by National Institutes of Health grants R35GM135160 (S.L.G.), R35GM130312 (J.E.B.) and American Heart Association Predoctoral Fellowship 906429 (C.T.H). Slide scanner imaging was performed at the UNC Hooker Imaging Core Facility, supported in part by P30 CA016086 Cancer Center Core Support Grant to the UNC Lineberger Comprehensive Cancer Center. Imaging was also supported by an award from the UNC Core Facilities Advocacy Committee and Office of Research Technologies, UNC Chapel Hill School of Medicine. Immunohistochemistry services were provided by the Histology Research Core Facility in the Department of Cell Biology and Physiology at the University of North Carolina, Chapel Hill NC (Ashley Ezzell).

## Author contributions

Conceptualization: C.T.H., J.E.B., S.L.G.; Methodology: C.T.H., E.B.E., K.L, E.C.O., B.T.; Formal analysis: C.T.H., E.B.E., K.L., E.C.O., B.T.; Investigation: C.T.H., E.B.E., K.L., A.S., C.-H.H; Resources: S.L.G.; Writing - original draft: C.T.H., S.L.G.; Writing - review & editing: C.T.H., E.B.E., K.L., E.C.O., A.S, C.-H.H., B.T., J.E.B., S.L.G.; Visualization: C.T.H., C.-H.H., S.L.G.; Supervision: S.L.G.; Project administration: S.L.G.; Funding acquisition: S.L.G.

## Supplementary Information

**Figure S1.**
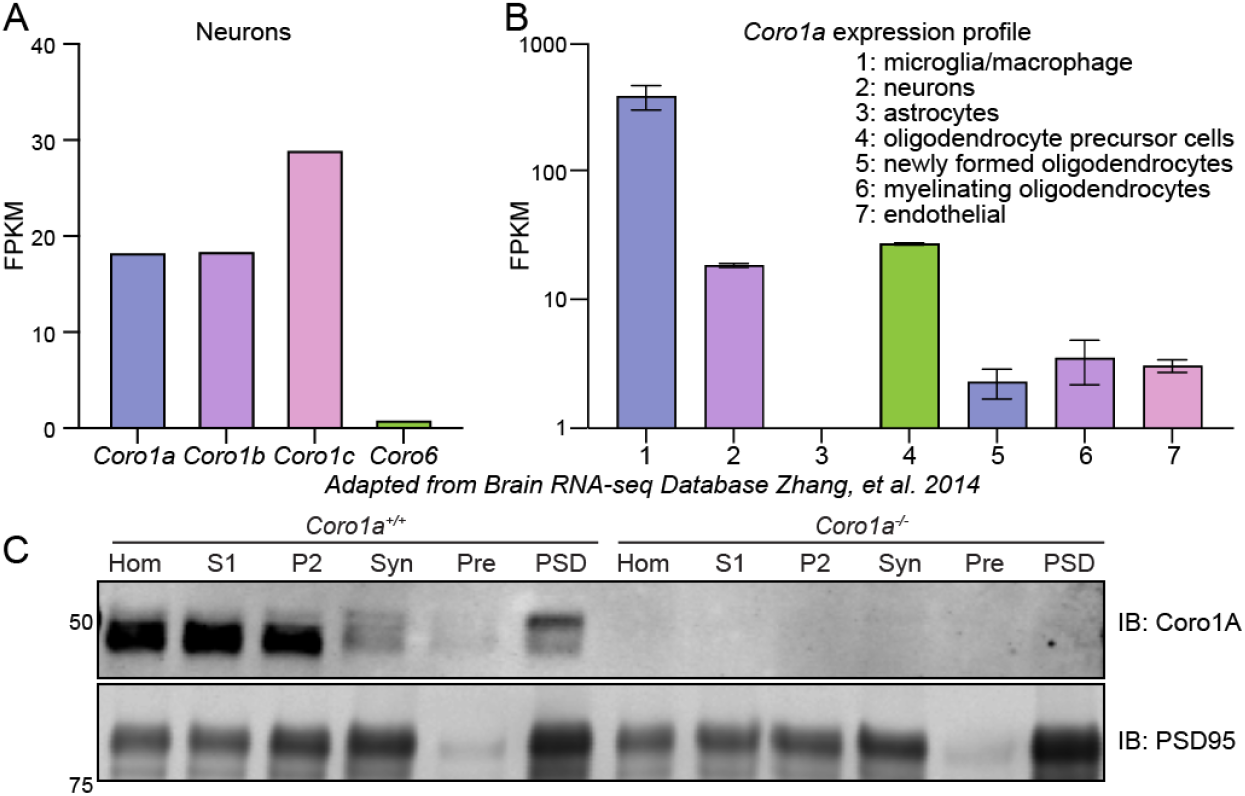
Coro1A is expressed in developing neurons and enriched in the postsynaptic density. (**A**) RNA-seq results adapted from the Brain RNA-seq Database (Zhang et al., 2014), demonstrating that Coro1A, Coro1B, and Coro1C are expressed in neurons. FPKM: Fragments Per Kilobase of transcript per Million mapped reads. (**B**) RNA-seq results adapted from the Brain RNA-seq Database, demonstrating that *Coro1a* expression profile across different brain cell types. (**C**) Western blotting of *Coro1a*+/+ and *Coro1a*-/-postsynaptic density fractionation samples. Hom (homogenate), S1 (Supernatant 1), P2 (Pellet 2), Syn (Synaptosome), Pre (Presynapse), PSD (Postsynaptic density).

**Figure S2.**
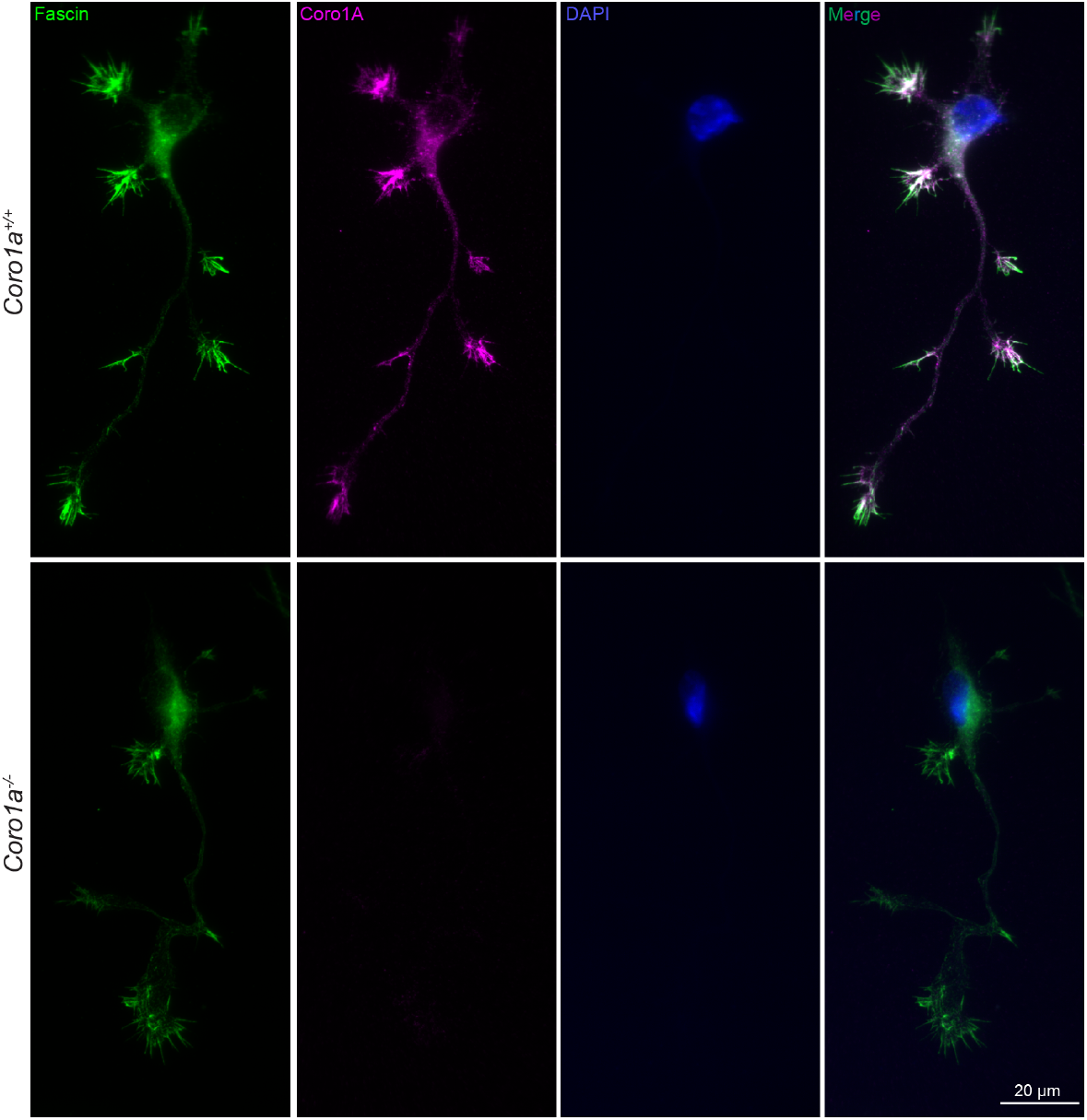
Validation of Coro1A antibody. Representative images of *Coro1a*+/+ or *Coro1a*-/-2 DIV neurons stained with Coro1A and fascin, showing the specificity of Coro1A antibody. Intensity for the same fluorescent channels were set to the same scale.

**Figure S3.**
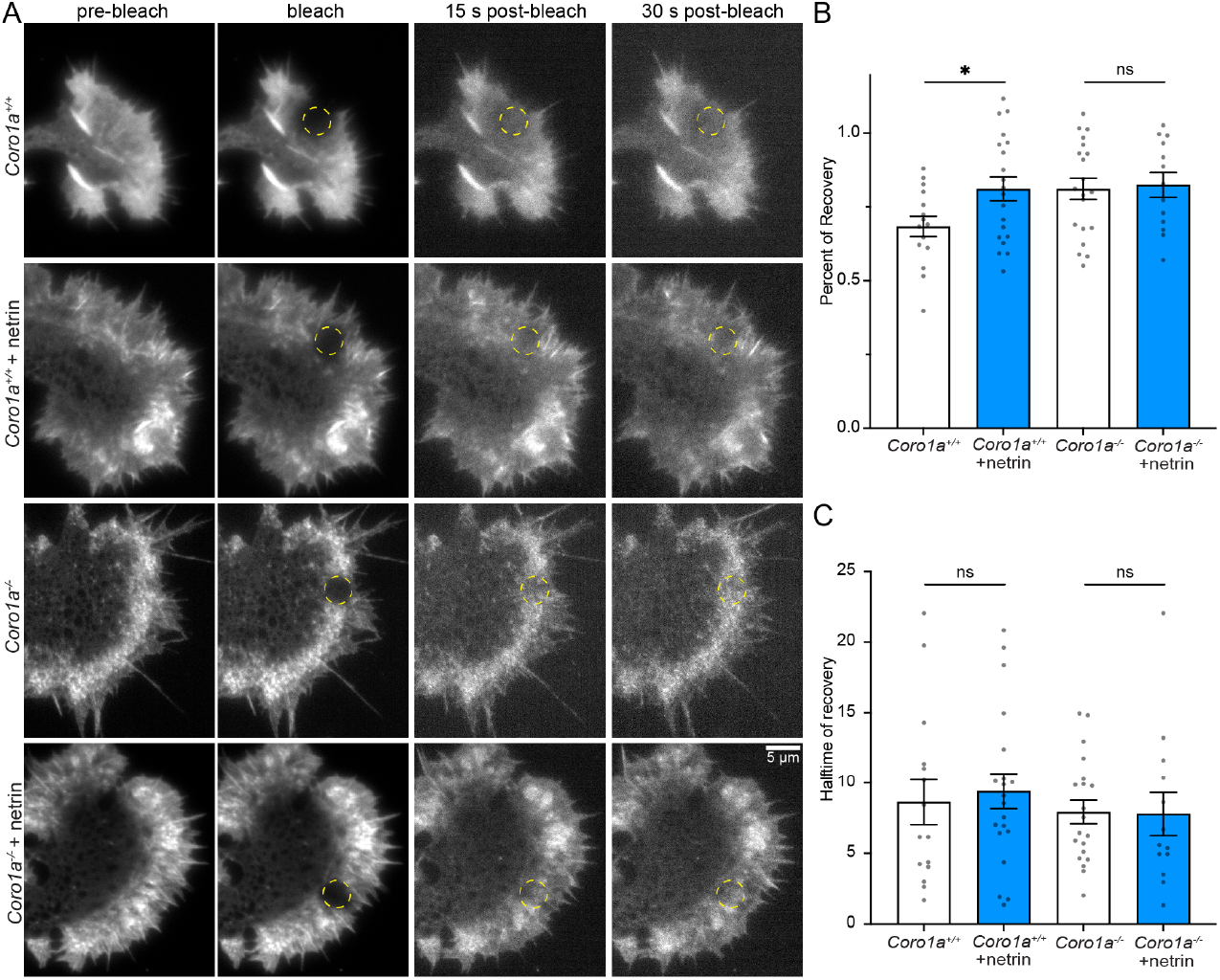
Coro1A is required for growth cone responses to netrin. (**A**) Representative images of the FRAP of GFP-actin in *Coro1a*+/+ or *Coro1a*-/-2 DIV neurons treated with or without netrin (600ng/mL) for 30 min. Yellow circle denotes the bleach ROI. For display purpose, each image was scaled based on minimum and maximum intensity. (**B**) Quantification of GFP-actin fluorescence recovery percentage (mobile fraction, plateau) from individual recovery curves. (**C**) Quantification of GFP-actin fluorescence recovery halftime (t1/2) from individual recovery curves. n(cells)= 15 *Coro1a*+/+, 20 *Coro1a*+/+ netrin, 20 *Coro1a*-/-media, 13 *Coro1a*-/-netrin from three biological replicates. Paired two-tailed t-test. *, p <0.05; ns, not statistically different.

**Figure S4.**
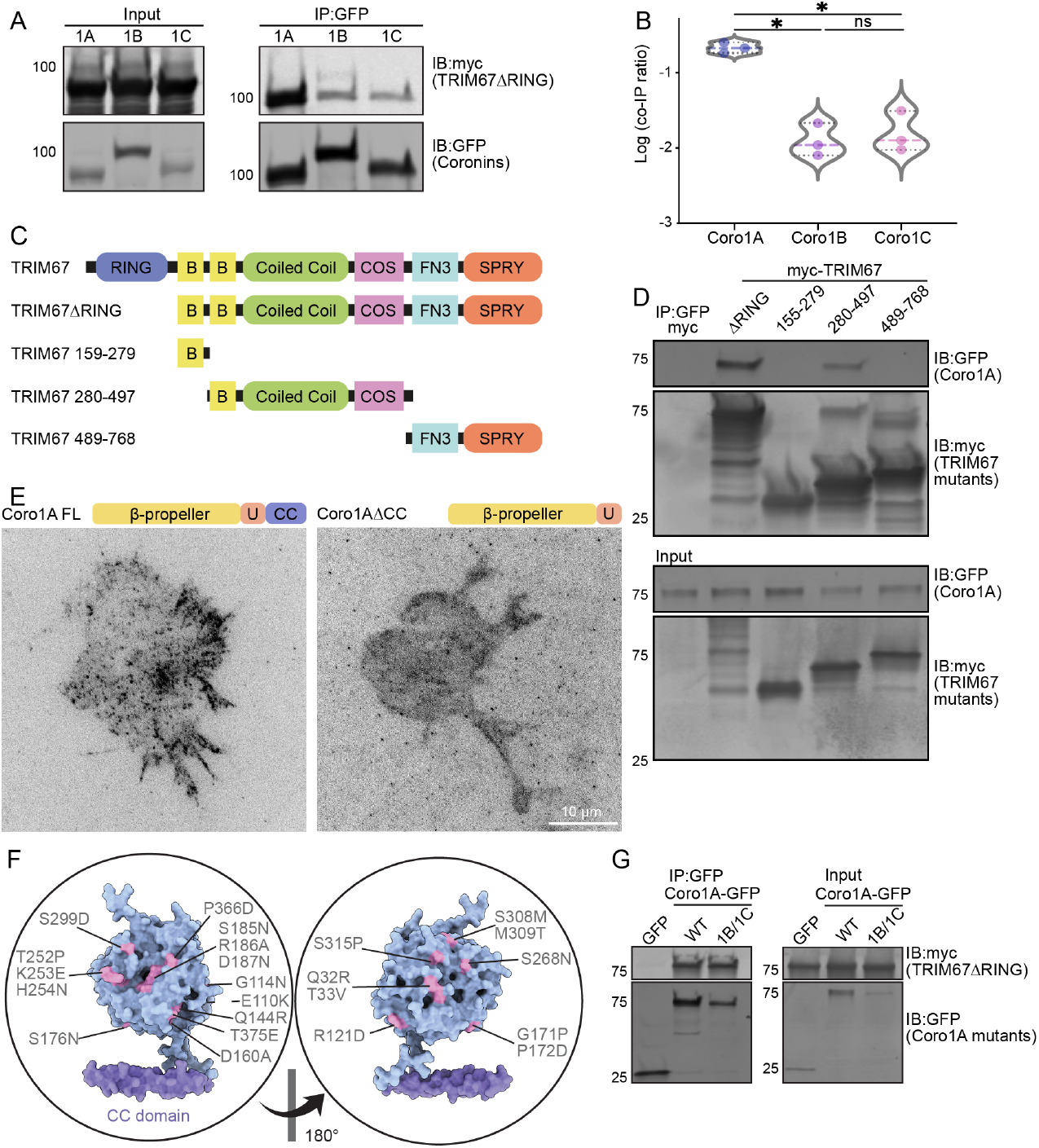
Mapping the interaction domains of TRIM67 and Coro1A. (**A**) Coimmunoprecipitation assay from HEK293 cells transfected with myc-TRIM67ΔRING and brain expressed coronins tagged with GFP demonstrating TRIM67ΔRING interacts with Coro1A, Coro1B, and Coro1C, with the interaction being strongest with Coro1A. (**B**) Quantification of myc-TRIM67ΔRING coimmunoprecipitation ratios from (A), showing or myc-TRIMΔRING levels rel-ative to GFP-tagged type-I coronins. n (cultures)= 3. One-way ANOVA (Dunnett’s multiple comparisons tests). *, p <0.05; ns, not statistically different. (**C**) Schematic of TRIM67 truncations used in structure-function assays. (**D**) Structure-function coimmuno-precipitation assays from *TRIM67^-/-^* HEK293 cells transfected with GFP-Coro1A and indicated myc-TRIM67ΔRING fragments, showing that fragment 280-497 (predicted dimerization region) that contains the coiled coil domain interacts with TRIM67. (**E**) Representative images from total internal reflection fluorescence (TIRF) live-cell imaging of 2 DIV cortical neurons expressing Coro1A-GFP or Coro1AΔCC-GFP, showing that loss of coiled coil domain alters Coro1A localization. (**F**) Coro1A monomer highlighting the Coro1A unique surface residues that were mutated in a single construct for mapping the Coro1A:TRIM67 inter-action. (**G**) Coimmunoprecipitation assays from HEK293 cells transfected with myc-TRIM67ΔRING and Coro1A-GFP or Coro1A β-propeller mutant, showing the unique surface residues on Coro1A β-propeller are not required for Coro1A:TRIM67 interaction.

## Notes

### Competing Interest Statement

The authors have declared no competing interest.

## Bibliography

Ahmed, Z., Shaw, G., Sharma, V. P., Yang, C., McGowan, E., and Dickson, D. W. (2007). Actin-binding proteins coronin-1a and IBA-1 are effective microglial markers for immunohistochemistry. The Journal of Histochemistry and Cytochemistry: Official Journal of the Histochemistry Society, 55(7):687–700. doi: 10.1369/jhc.6A7156.2007. PMID: 17341475.

Appleton, B. A., Wu, P., and Wiesmann, C. (2006). The crystal structure of murine coronin-1: a regulator of actin cytoskeletal dynamics in lymphocytes. Structure (London, England : 1993), 14(1):87–96. doi: 10.1016/j.str.2005.09.013. PMID: 16407068 publisher-place: United States.

Asano, S., Mishima, M., and Nishida, E. (2001). Coronin forms a stable dimer through its C-terminal coiled coil region: an implicated role in its localization to cell periphery. Genes to Cells, 6(3):225–235. doi: 10.1046/j.1365-2443.2001.00416.x. _eprint: https://onlinelibrary.wiley.com/doi/pdf/10.1046/j.1365-2443.2001.00416.x.

Askew, K. and Gomez-Nicola, D. (2018). A story of birth and death: Insights into the formation and dynamics of the microglial population. Brain, Behavior, and Immunity, 69:9–17. doi: 10.1016/j.bbi.2017.03.009.

Bastmeyer, M. and O’Leary, D. D. (1996). Dynamics of target recognition by interstitial axon branching along developing cortical axons. The Journal of Neuroscience: The Official Journal of the Society for Neuroscience, 16(4):1450–1459. doi: 10.1523/JNEUROSCI.16-04-01450.1996. PMID: 8778296 PMCID: PMC6578553.

Bin, J. M., Han, D., Lai Wing Sun, K., Croteau, L.-P., Dumontier, E., Cloutier, J.-F., Kania, A., and Kennedy, T. E. (2015). Complete Loss of Netrin-1 Results in Embryonic Lethality and Severe Axon Guidance Defects without Increased Neural Cell Death. Cell Reports, 12(7):1099–1106. doi: 10.1016/j.celrep.2015.07.028.

Blaker-Lee, A., Gupta, S., McCammon, J. M., De Rienzo, G., and Sive, H. (2012). Zebrafish homologs of genes within 16p11.2, a genomic region associated with brain disorders, are active during brain development, and include two deletion dosage sensor genes. Disease models & mechanisms, 5(6):834–851. doi: 10.1242/dmm.009944. PMID: 22566537.

Bodmer, D. and Kuruvilla, R. Trafficking of Trk Receptors, pages 273–289. Humana Press, Totowa, NJ, (2012). ISBN 978-1-61779-824-5. URL https://doi.org/10.1007/978-1-61779-824-5_16. DOI: 10.1007/978-1-61779-824-5_16.

Borovac, J., Bosch, M., and Okamoto, K. (2018). Regulation of actin dynamics during structural plasticity of dendritic spines: Signaling messengers and actin-binding proteins. Molecular and Cellular Neuroscience, 91:122–130. doi: 10.1016/j.mcn.2018.07.001.

Boyer, N. P. and Gupton, S. L. (2018). Revisiting Netrin-1: One Who Guides (Axons). Frontiers in Cellular Neuroscience, 12:221.

Boyer, N. P., Monkiewicz, C., Menon, S., Moy, S. S., and Gupton, S. L. (2018). Mammalian TRIM67 Functions in Brain Development and Behavior. eneuro, 5(3):ENEURO.0186– 18.2018. doi: 10.1523/ENEURO.0186-18.2018.

Boyer, N. P., McCormick, L. E., Menon, S., Urbina, F. L., and Gupton, S. L. (2020). A pair of E3 ubiquitin ligases compete to regulate filopodial dynamics and axon guidance. Journal of Cell Biology, 219(1). doi: 10.1083/jcb.201902088.

Brieher, W. M., Kueh, H. Y., Ballif, B. A., and Mitchison, T. J. (2006). Rapid actin monomer– insensitive depolymerization of Listeria actin comet tails by cofilin, coronin, and Aip1. Journal of Cell Biology, 175(2):315–324. doi: 10.1083/jcb.200603149.

Cai, L., Holoweckyj, N., Schaller, M. D., and Bear, J. E. (2005). Phosphorylation of coronin 1b by protein kinase C regulates interaction with Arp2/3 and cell motility. Journal of Biological Chemistry, 280(36):31913–31923. doi: 10.1074/jbc.M504146200. PMID: 16027158 publisher: Elsevier.

Cai, L., Makhov, A. M., and Bear, J. E. (2007). F-actin binding is essential for coronin 1b function in vivo. Journal of Cell Science, 120(10):1779 LP – 1790. doi: 10.1242/jcs.007641.

Cai, L., Marshall, T. W., Uetrecht, A. C., Schafer, D. A., and Bear, J. E. (2007). Coronin 1b coordinates Arp2/3 complex and cofilin activities at the leading edge. Cell, 128(5):915–929. doi: 10.1016/j.cell.2007.01.031.

Cai, L., Makhov, A. M., Schafer, D. A., and Bear, J. E. (2008). Coronin 1b antagonizes cortactin and remodels Arp2/3-containing actin branches in lamellipodia. Cell, 134(5):828–842. doi: 10.1016/j.cell.2008.06.054. PMID: 18775315.

Chan, K. T., Creed, S. J., and Bear, J. E. (2011). Unraveling the enigma: progress towards understanding the coronin family of actin regulators. Trends in cell biology, 21(8):481– 488. doi: 10.1016/j.tcb.2011.04.004.

Chan, K. T., Roadcap, D. W., Holoweckyj, N., and Bear, J. E. (2012). Coronin 1c harbours a second actin-binding site that confers co-operative binding to F-actin. The Biochemical journal, 444(1):89–96. doi: 10.1042/BJ20120209. PMID: 22364218 publisher: Portland Press Ltd.

Chen, Y., Ip, F. C. F., Shi, L., Zhang, Z., Tang, H., Ng, Y. P., Ye, W.-C., Fu, A. K. Y., and Ip, N. Y. (2014). Coronin 6 Regulates Acetylcholine Receptor Clustering through Modulating Receptor Anchorage to Actin Cytoskeleton. The Journal of Neuroscience, 34(7):2413. doi: 10.1523/JNEUROSCI.3226-13.2014.

de Hostos, E. L., Bradtke, B., Lottspeich, F., Guggenheim, R., and Gerisch, G. (1991). Coronin, an actin binding protein of Dictyostelium discoideum localized to cell surface projections, has sequence similarities to G protein beta subunits. The EMBO journal, 10 (13):4097–4104. PMID: 1661669.

Dent, E. W. and Gertler, F. B. (2003). Cytoskeletal dynamics and transport in growth cone motility and axon guidance. Neuron, 40(2):209–227. doi: 10.1016/s0896-6273(03)00633-0. PMID: 14556705 publisher-place: United States.

Dent, E. W., Barnes, A. M., Tang, F., and Kalil, K. (2004). Netrin-1 and Semaphorin 3a Promote or Inhibit Cortical Axon Branching, Respectively, by Reorganization of the Cytoskeleton. Journal of Neuroscience, 24(12):3002–3012. doi: 10.1523/JNEUROSCI.4963-03.2004. publisher: Society for Neuroscience section: Development/Plasticity/Repair PMID: 15044539.

Dent, E. W., Gupton, S. L., and Gertler, F. B. (2011). The growth cone cytoskeleton in axon outgrowth and guidance. Cold Spring Harbor perspectives in biology, 3(3). doi: 10.1101/cshperspect.a001800. PMID: 21106647.

Deshaies, R. J. and Joazeiro, C. A. P. (2009). Ring domain E3 ubiquitin ligases. Annual review of biochemistry, 78:399–434. doi: 10.1146/annurev.biochem.78.101807.093809. PMID: 19489725 publisher-place: United States.

Dudley-Fraser, J. and Rittinger, K. (2023). It’s a TRIM-endous view from the top: the varied roles of TRIpartite Motif proteins in brain development and disease. Frontiers in Molecular Neuroscience, 16. doi: 10.3389/fnmol.2023.1287257. publisher: Frontiers.

Escamilla, C. O., Filonova, I., Walker, A. K., Xuan, Z. X., Holehonnur, R., Espinosa, F., Liu, S., Thyme, S. B., López-García, I. A., Mendoza, D. B., Usui, N., Ellegood, J., Eisch, A. J., Konopka, G., Lerch, J. P., Schier, A. F., Speed, H. E., and Powell, C. M. (2017). Kctd13 deletion reduces synaptic transmission via increased RhoA. Nature, 551(7679):227–231. doi: 10.1038/nature24470. publisher: Nature Publishing Group.

Evans, R., O’Neill, M., Pritzel, A., Antropova, N., Senior, A., Green, T., Žídek, A., Bates, R., Blackwell, S., Yim, J., Ronneberger, O., Bodenstein, S., Zielinski, M., Bridgland, A., Potapenko, A., Cowie, A., Tunyasuvunakool, K., Jain, R., Clancy, E., Kohli, P., Jumper, J., and Hassabis, D. (2022). Protein complex prediction with AlphaFold-Multimer. doi: 10.1101/2021.10.04.463034. page: 2021.10.04.463034 section: New Results.

Fazeli, A., Dickinson, S. L., Hermiston, M. L., Tighe, R. V., Steen, R. G., Small, C. G., Stoeckli, E. T., Keino-Masu, K., Masu, M., Rayburn, H., Simons, J., Bronson, R. T., Gordon, J. I., Tessier-Lavigne, M., and Weinberg, R. A. (1997). Phenotype of mice lacking functional Deleted in colorectal cancer (Dec) gene. Nature, 386(6627):796–804. doi: 10.1038/386796a0. publisher: Nature Publishing Group.

Flores, C. (2011). Role of netrin-1 in the organization and function of the mesocorticolimbic dopamine system. Journal of psychiatry & neuroscience : JPN, 36(5):296–310. doi: 10.1503/jpn.100171. PMID: 21481303 publisher: Canadian Medical Association.

Föger, N., Rangell, L., Danilenko, D. M., and Chan, A. C. (2006). Requirement for Coronin 1 in T Lymphocyte Trafficking and Cellular Homeostasis. Science, 313(5788):839 LP – 842. doi: 10.1126/science.1130563.

Gallo, G. (2011). The cytoskeletal and signaling mechanisms of axon collateral branching. Developmental Neurobiology, 71(3):201–220. doi: 10.1002/dneu.20852. PMID: 21308993.

Gandhi, M., Achard, V., Blanchoin, L., and Goode, B. L. (2009). Coronin Switches Roles in Actin Disassembly Depending on the Nucleotide State of Actin. Molecular Cell, 34(3):364–374. doi: 10.1016/j.molcel.2009.02.029.

Gatfield, J., Albrecht, I., Zanolari, B., Steinmetz, M. O., and Pieters, J. (2005). Association of the leukocyte plasma membrane with the actin cytoskeleton through coiled coil-mediated trimeric coronin 1 molecules. Molecular Biology of the Cell, 16(6):2786–2798. doi: 10.1091/mbc.e05-01-0042. PMID: 15800061 PMCID: PMC1142424.

Gay, S. M., Chartampila, E., Lord, J. S., Grizzard, S., Maisashvili, T., Ye, M., Barker, N. K., Mordant, A. L., Mills, C. A., Herring, L. E., and Diering, G. H. (2024). Developing forebrain synapses are uniquely vulnerable to sleep loss. Proceedings of the National Academy of Sciences, 121(44):e2407533121. doi: 10.1073/pnas.2407533121. publisher: Proceedings of the National Academy of Sciences.

Glasgow, S. D., Labrecque, S., Beamish, I. V., Aufmkolk, S., Gibon, J., Han, D., Harris, S. N., Dufresne, P., Wiseman, P. W., McKinney, R. A., Séguéla, P., De Koninck, P., Ruthazer, E. S., and Kennedy, T. E. (2018). Activity-Dependent Netrin-1 Secretion Drives Synaptic Insertion of GluA1-Containing AMPA Receptors in the Hippocampus. Cell Reports, 25 (1):168–182.e6. doi: 10.1016/j.celrep.2018.09.028.

Goldman, J. S., Ashour, M. A., Magdesian, M. H., Tritsch, N. X., Harris, S. N., Christofi, N., Chemali, R., Stern, Y. E., Thompson-Steckel, G., Gris, P., Glasgow, S. D., Grutter, P., Bouchard, J.-F., Ruthazer, E. S., Stellwagen, D., and Kennedy, T. E. (2013). Netrin-1 Promotes Excitatory Synaptogenesis between Cortical Neurons by Initiating Synapse Assembly. Journal of Neuroscience, 33(44):17278–17289. doi: 10.1523/JNEUROSCI.1085-13.2013. publisher: Society for Neuroscience section: Articles PMID: 24174661.

Goode, B. L., Wong, J. J., Butty, A. C., Peter, M., McCormack, A. L., Yates, J. R., Drubin, D. G., and Barnes, G. (1999). Coronin promotes the rapid assembly and cross-linking of actin filaments and may link the actin and microtubule cytoskeletons in yeast. The Journal of cell biology, 144(1):83–98. doi: 10.1083/jcb.144.1.83. PMID: 9885246 publisher: The Rockefeller University Press.

Grogan, A., Reeves, E., Keep, N., Wientjes, F., Totty, N. F., Burlingame, A. L., Hsuan, J. J., and Segal, A. W. (1997). Cytosolic phox proteins interact with and regulate the assembly of coronin in neutrophils. Journal of Cell Science, 110 (Pt 24):3071–3081. doi: 10.1242/jcs.110.24.3071. PMID: 9365277.

Han, X., Hu, Z., Surya, W., Ma, Q., Zhou, F., Nordenskiöld, L., Torres, J., Lu, L., and Miao, Y. (2023). The intrinsically disordered region of coronins fine-tunes oligomerization and actin polymerization. Cell Reports, 42(6):112594. doi: 10.1016/j.celrep.2023.112594. PMID: 37269287.

Hao, J. C., Adler, C. E., Mebane, L., Gertler, F. B., Bargmann, C. I., and Tessier-Lavigne, M. (2010). The Tripartite Motif Protein MADD-2 Functions with the Receptor UNC-40 (DCC) in Netrin-Mediated Axon Attraction and Branching. Developmental Cell, 18(6):950–960. doi: 10.1016/j.devcel.2010.02.019.

Ho, C. T. and Gupton, S. L. Neurons | Establishing and Maintaining Neuron Morphology, pages 345–357. Elsevier, Oxford, (2021). ISBN 978-0-12-822040-5. URL https://www.sciencedirect.com/science/article/pii/B9780128194607002401. DOI 10.1016/B978-0-12-819460-7.00240-1.

Hotulainen, P. and Hoogenraad, C. C. (2010). Actin in dendritic spines: connecting dynamics to function. Journal of Cell Biology, 189(4):619–629. doi: 10.1083/jcb.201003008.

Hoyer, M. J., Chitwood, P. J., Ebmeier, C. C., Striepen, J. F., Qi, R. Z., Old, W. M., and Voeltz, G. K. (2018). A Novel Class of ER Membrane Proteins Regulates ER-Associated Endosome Fission. Cell, 175(1):254–265.e14. doi: 10.1016/j.cell.2018.08.030.

Humphries, C. L., Balcer, H. I., D’Agostino, J. L., Winsor, B., Drubin, D. G., Barnes, G., Andrews, B. J., and Goode, B. L. (2002). Direct regulation of Arp2/3 complex activity and function by the actin binding protein coronin. The Journal of cell biology, 159(6):993–1004. doi: 10.1083/jcb.200206113. PMID: 12499356.

Hylton, R. K., Heebner, J. E., Grillo, M. A., and Swulius, M. T. (2022). Cofilactin filaments regulate filopodial structure and dynamics in neuronal growth cones. Nature Communications, 13(1):2439. doi: 10.1038/s41467-022-30116-x. PMID: 35508487 PMCID: PMC9068697.

Iconomou, M. and Saunders, D. N. (2016). Systematic approaches to identify E3 ligase substrates. Biochemical Journal, 473(22):4083–4101. doi: 10.1042/BCJ20160719.

Jacquemet, G. Mapping the Localization of Proteins Within Filopodia Using FiloMap, pages 51–61. Springer US, New York, NY, (2023). ISBN 978-1-07-162887-4. URL https://doi.org/10.1007/978-1-0716-2887-4_4. DOI: 10.1007/978-1-0716-2887-4_4.

Jacquemet, G., Stubb, A., Saup, R., Miihkinen, M., Kremneva, E., Hamidi, H., and Ivaska, J. (2019). Filopodome Mapping Identifies p130cas as a Mechanosensitive Regulator of Filopodia Stability. Current biology: CB, 29(2):202–216.e7. doi: 10.1016/j.cub.2018.11.053. PMID: 30639111 PMCID: PMC6345628.

Jansen, S., Collins, A., Chin, S. M., Ydenberg, C. A., Gelles, J., and Goode, B. L. (2015). Single-molecule imaging of a three-component ordered actin disassembly mechanism. Nature Communications, 6(1):7202. doi: 10.1038/ncomms8202.

Jayachandran, R., Liu, X., BoseDasgupta, S., Müller, P., Zhang, C.-L., Moshous, D., Studer, V., Schneider, J., Genoud, C., Fossoud, C., Gambino, F., Khelfaoui, M., Müller, C., Bartholdi, D., Rossez, H., Stiess, M., Houbaert, X., Jaussi, R., Frey, D., Kammerer, R. A., Deupi, X., de Villartay, J.-P., Lüthi, A., Humeau, Y., and Pieters, J. (2014). Coronin 1 Regulates Cognition and Behavior through Modulation of cAMP/Protein Kinase A Signaling. PLOS Biology, 12(3):e1001820. publisher: Public Library of Science.

Jumper, J., Evans, R., Pritzel, A., Green, T., Figurnov, M., Ronneberger, O., Tunyasuvunakool, K., Bates, R., Žídek, A., Potapenko, A., Bridgland, A., Meyer, C., Kohl, S. A. A., Ballard, A. J., Cowie, A., Romera-Paredes, B., Nikolov, S., Jain, R., Adler, J., Back, T., Petersen, S., Reiman, D., Clancy, E., Zielinski, M., Steinegger, M., Pacholska, M., Berghammer, T., Bodenstein, S., Silver, D., Vinyals, O., Senior, A. W., Kavukcuoglu, K., Kohli, P., and Hassabis, D. (2021). Highly accurate protein structure prediction with AlphaFold. Nature, 596(7873):583–589. doi: 10.1038/s41586-021-03819-2. number: 7873 publisher: Nature Publishing Group.

Kalil, K. and Dent, E. W. (2014). Branch management: mechanisms of axon branching in the developing vertebrate CNS. Nature reviews. Neuroscience, 15(1):7–18. doi: 10.1038/nrn3650. PMID: 24356070.

Kammerer, R. A., Kostrewa, D., Progias, P., Honnappa, S., Avila, D., Lustig, A., Winkler, F. K., Pieters, J., and Steinmetz, M. O. (2005). A conserved trimerization motif controls the topology of short coiled coils. Proceedings of the National Academy of Sciences of the United States of America, 102(39):13891–13896. doi: 10.1073/pnas.0502390102. PMID: 16172398 PMCID: PMC1236532.

Ketschek, A. and Gallo, G. (2010). Nerve growth factor induces axonal filopodia through localized microdomains of phosphoinositide 3-kinase activity that drive the formation of cytoskeletal precursors to filopodia. The Journal of neuroscience : the official journal of the Society for Neuroscience, 30(36):12185–12197. doi: 10.1523/JNEUROSCI.1740-10.2010. publisher: Society for Neuroscience.

Kim, T. Y., Siesser, P. F., Rossman, K. L., Goldfarb, D., Mackinnon, K., Yan, F., Yi, X., MacCoss, M. J., Moon, R. T., Der, C. J., and Major, M. B. (2015). Substrate trapping proteomics reveals targets of the βTrCP2/FBXW11 ubiquitin ligase. Molecular and cellular biology, 35(1):167–181. doi: 10.1128/MCB.00857-14. publisher: American Society for Microbiology.

King, Z. T., Butler, M. T., Hockenberry, M. A., Subramanian, B. C., Siesser, P. F., Graham, D. M., Legant, W. R., and Bear, J. E. (2022). Coro1b and Coro1C regulate lamellipodia dynamics and cell motility by tuning branched actin turnover. Journal of Cell Biology, 221 (8):e202111126. doi: 10.1083/jcb.202111126.

Kizner, V., Naujock, M., Fischer, S., Jäger, S., Reich, S., Schlotthauer, I., Zuckschwerdt, K., Geiger, T., Hildebrandt, T., Lawless, N., Macartney, T., Dorner-Ciossek, C., and Gillardon, F. (2020). Crispr/Cas9-mediated Knockout of the Neuropsychiatric Risk Gene KCTD13 Causes Developmental Deficits in Human Cortical Neurons Derived from Induced Pluripotent Stem Cells. Molecular Neurobiology, 57(2):616–634. doi:10.1007/s12035-019-01727-1. PMID: 31402430.

Kolodkin, A. L. and Tessier-Lavigne, M. (2011). Mechanisms and molecules of neuronal wiring: a primer. Cold Spring Harbor perspectives in biology, 3(6):a001727. doi: 10.1101/cshperspect.a001727. publisher: Cold Spring Harbor Laboratory Press.

Konietzny, A., Bär, J., and Mikhaylova, M. (2017). Dendritic Actin Cytoskeleton: Structure, Functions, and Regulations. Frontiers in Cellular Neuroscience, 11. doi: 10.3389/fncel.2017.00147. publisher: Frontiers.

Korobova, F. and Svitkina, T. (2008). Arp2/3 Complex Is Important for Filopodia Formation, Growth Cone Motility, and Neuritogenesis in Neuronal Cells. Molecular Biology of the Cell, 19(4):1561–1574. doi: 10.1091/mbc.e07-09-0964. publisher: American Society for Cell Biology (mboc).

Koundinya, N., Aguilar, R. M., Wetzel, K., Tomasso, M. R., Nagarajan, P., McGuirk, E. R., Padrick, S. B., and Goode, B. L. (2025). Two ligands of Arp2/3 complex, yeast coronin and GMF, interact and synergize in pruning branched actin networks. Journal of Biological Chemistry, 301(3):108191. doi: 10.1016/j.jbc.2025.108191.

Lewis, A. K. and Bridgman, P. C. (1992). Nerve growth cone lamellipodia contain two populations of actin filaments that differ in organization and polarity. The Journal of Cell Biology, 119(5):1219–1243. doi: 10.1083/jcb.119.5.1219. PMID: 1447299 PMCID: PMC2289720.

Lim, K. L., Chew, K. C. M., Tan, J. M. M., Wang, C., Chung, K. K. K., Zhang, Y., Tanaka, Y., Smith, W., Engelender, S., Ross, C. A., Dawson, V. L., and Dawson, T. M. (2005). Parkin Mediates Nonclassical, Proteasomal-Independent Ubiquitination of Synphilin-1: Implications for Lewy Body Formation. Journal of Neuroscience, 25(8):2002–2009. doi: 10.1523/JNEUROSCI.4474-04.2005. publisher: Society for Neuroscience section: Neurobiology of Disease PMID: 15728840.

Liu, S.-L., Needham, K. M., May, J. R., and Nolen, B. J. (2011). Mechanism of a Concentration-dependent Switch between Activation and Inhibition of Arp2/3 Complex by Coronin*. Journal of Biological Chemistry, 286(19):17039–17046. doi: 10.1074/jbc.M111.219964.

Longair, M. H., Baker, D. A., and Armstrong, J. D. (2011). Simple Neurite Tracer: open source software for reconstruction, visualization and analysis of neuronal processes. Bioinformatics (Oxford, England), 27(17):2453–2454. doi: 10.1093/bioinformatics/btr390. PMID: 21727141.

Loo, L., Simon, J. M., Xing, L., McCoy, E. S., Niehaus, J. K., Guo, J., Anton, E. S., and Zylka, M. J. (2019). Single-cell transcriptomic analysis of mouse neocortical development. Nature Communications, 10(1):134. doi: 10.1038/s41467-018-08079-9. PMID: 30635555 PMCID: PMC6329831.

Lowery, L. A. and Vactor, D. V. (2009). The trip of the tip: Understanding the growth cone machinery. Nature Reviews Molecular Cell Biology, 10(5):332–343. doi: 10.1038/nrm2679.

Machesky, L. M., Reeves, E., Wientjes, F., Mattheyse, F. J., Grogan, A., Totty, N. F., Burlingame, A. L., Hsuan, J. J., and Segal, A. W. (1997). Mammalian actin-related protein 2/3 complex localizes to regions of lamellipodial protrusion and is composed of evolutionarily conserved proteins. The Biochemical Journal, 328 (Pt 1)(Pt 1):105–112. doi: 10.1042/bj3280105. PMID: 9359840 PMCID: PMC1218893.

McArdle, B. and Hofmann, A. (2008). Coronin structure and implications. Sub-cellular biochemistry, 48:56–71. doi: 10.1007/978-0-387-09595-0_6. PMID: 18925371 publisher-place: United States.

McCarthy, S., Makarov, V., Kirov, G., Addington, A., McClellan, J., Yoon, S., Perkins, D., Dickel, D. E., Kusenda, M., Krastoshevsky, O., Krause, V., Kumar, R. A., Grozeva, D., Malhotra, D., Walsh, T., Zackai, E. H., Kaplan, P., Ganesh, J., Krantz, I. D., Spinner, N. B., Roccanova, P., Bhandari, A., Pavon, K., Lakshmi, B., Leotta, A., Kendall, J., Lee, Y.-h., Vacic, V., Gary, S., Iakoucheva, L., Crow, T. J., Christian, S. L., Lieberman, J., Stroup, S., Lehtimäki, T., Puura, K., Haldeman-Englert, C., Pearl, J., Goodell, M., Willour, V. L., DeRosse, P., Steele, J., Kassem, L., Wolff, J., Chitkara, N., McMahon, F. J., Malhotra, A. K., Potash, J. B., Schulze, T. G., Nöthen, M. M., Cichon, S., Rietschel, M., Leibenluft, E., Kustanovich, V., Lajonchere, C. M., Sutcliffe, J. S., Skuse, D., Gill, M., Gallagher, L., Mendell, N. R., Craddock, N., Owen, M. J., O’Donovan, M. C., Shaikh, T. H., Susser, E., DeLisi, L. E., Sullivan, P. F., Deutsch, C. K., Rapoport, J., Levy, D. L., King, M.-C., and Sebat, J. (2009). Microduplications of 16p11.2 are Associated with Schizophrenia. Nature genetics, 41(11):1223–1227. doi: 10.1038/ng.474. PMID: 19855392 PMCID:PMC2951180.

McCormick, L. E. and Gupton, S. L. (2020). Mechanistic advances in axon pathfinding. Current Opinion in Cell Biology, 63:11–19. doi: 10.1016/j.ceb.2019.12.003.

Mccormick, L. E., Baker, N. K., Herring, L. E., and Gupton, S. L. (2024). Loss of the E3 ubiquitin ligase TRIM67 alters the post-synaptic density proteome. microPublication Biology. doi: 10.17912/micropub.biology.001118. publisher: Caltech Library.

McCormick, L. E., Evans, E. B., Barker, N. K., Herring, L. E., Diering, G. H., and Gupton, S. L. (2024). The E3 ubiquitin ligase TRIM9 regulates synaptic function and actin dynamics in response to netrin-1. Molecular Biology of the Cell, 35(5):ar67. doi: 10.1091/mbc.E23-12-0476. publisher: American Society for Cell Biology (mboc).

McCormick, L. E., Suarez, C., Herring, L. E., Cannon, K. S., Kovar, D. R., Brown, N. G., and Gupton, S. L. (2024). Multi-monoubiquitylation controls VASP-mediated actin dynamics. Journal of Cell Science, 137(2):jcs261527. doi: 10.1242/jcs.261527. PMID: 38277158 PMCID: PMC10917064.

Menon, S., Boyer, N. P., Winkle, C. C., McClain, L. M., Hanlin, C. C., Pandey, D., Rothenfußer, S., Taylor, A. M., and Gupton, S. L. (2015). The E3 Ubiquitin Ligase TRIM9 Is a Filopodia Off Switch Required for Netrin-Dependent Axon Guidance. Developmental cell, 35(6):698–712. doi: 10.1016/j.devcel.2015.11.022.

Menon, S., Goldfarb, D., Ho, C. T., Cloer, E. W., Boyer, N. P., Hardie, C., Bock, A. J., Johnson, E. C., Anil, J., Major, M. B., and Gupton, S. L. (2021). The TRIM9/TRIM67 neuronal interactome reveals novel activators of morphogenesis. Molecular biology of the cell, 32 (4):314–330. doi: 10.1091/mbc.E20-10-0622. PMID: 33378226 publisher-place: United States.

Meroni, G. and Diez-Roux, G. (2005). Trim/RBCC, a novel class of ‘single protein RING finger’ E3 ubiquitin ligases. BioEssays : news and reviews in molecular, cellular and developmental biology, 27(11):1147–1157. doi: 10.1002/bies.20304. PMID: 16237670 publisher-place: United States.

Minamide, L. S., Hylton, R., Swulius, M., and Bamburg, J. R. (2023). Visualizing Cofilin-Actin Filaments by Immunofluorescence and CryoEM: Essential Steps for Observing Cofilactin in Cells. Methods in Molecular Biology (Clifton, N.J.), 2593:265–281. doi: 10.1007/978-1-0716-2811-9_18. PMID: 36513938.

Mishima, M. and Nishida, E. (1999). Coronin localizes to leading edges and is involved in cell spreading and lamellipodium extension in vertebrate cells. Journal of Cell Science, 112(17):2833–2842. doi: 10.1242/jcs.112.17.2833.

Morikawa, R. K., Kanamori, T., Yasunaga, K.-i., and Emoto, K. (2011). Different levels of the Tripartite motif protein, Anomalies in sensory axon patterning (Asap), regulate distinct axonal projections of Drosophila sensory neurons. Proceedings of the National Academy of Sciences, 108(48):19389–19394. doi: 10.1073/pnas.1109843108. publisher: Proceedings of the National Academy of Sciences.

Mutalik, S. P., Ho, C. T., O’Shaughnessy, E. C., Frasineanu, A. G., Shah, A. B., and Gupton, S. L. (2025). Trim9 Controls Growth Cone Responses to Netrin Through DCC and UNC5C. Journal of Neurochemistry, 169(1):e70002. doi: 10.1111/jnc.70002._eprint: https://onlinelibrary.wiley.com/doi/pdf/10.1111/jnc.70002.

Nal, B., Carroll, P., Mohr, E., Verthuy, C., Da Silva, M.-I., Gayet, O., Guo, X.-J., He, H.-T., Alcover, A., and Ferrier, P. (2004). Coronin-1 expression in T lymphocytes: insights into protein function during T cell development and activation. International immunology, 16(2):231–240. doi: 10.1093/intimm/dxh022. PMID: 14734608 publisher-place: England.

Niarchou, M., Chawner, S. J. R. A., Doherty, J. L., Maillard, A. M., Jacquemont, S., Chung, W. K., Green-Snyder, L., Bernier, R. A., Goin-Kochel, R. P., Hanson, E., Linden, D. E. J., Linden, S. C., Raymond, F. L., Skuse, D., Hall, J., Owen, M. J., and Bree, M. B. M. (2019). Psychiatric disorders in children with 16p11.2 deletion and duplication. Translational Psychiatry, 9(1):1–8. doi: 10.1038/s41398-018-0339-8. publisher: Nature Publishing Group.

Nikodemova, M., Kimyon, R. S., De, I., Small, A. L., Collier, L. S., and Watters, J. J. (2015). Microglial numbers attain adult levels after undergoing a rapid decrease in cell number in the third postnatal week. Journal of Neuroimmunology, 278:280–288. doi: 10.1016/j.jneuroim.2014.11.018. PMID: 25468773 PMCID: PMC4297717.

Plooster, M., Menon, S., Winkle, C. C., Urbina, F. L., Monkiewicz, C., Phend, K. D., Weinberg, R. J., and Gupton, S. L. (2017). Trim9-dependent ubiquitination of DCC constrains kinase signaling, exocytosis, and axon branching. Molecular Biology of the Cell, 28(18):2374– 2385. doi: 10.1091/mbc.E16-08-0594. PMID: 28701345 PMCID: PMC5576901.

Pollard, T. D. and Cooper, J. A. (2009). Actin, a central player in cell shape and movement. Science (New York, N.Y.), 326(5957):1208–1212. doi: 10.1126/science.1175862. PMID: 19965462.

Popović, A., Miihkinen, M., Ghimire, S., Saup, R., Grönloh, M. L. B., Ball, N. J., Goult, B. T., Ivaska, J., and Jacquemet, G. (2023). Myosin-X recruits lamellipodin to filopodia tips. Journal of Cell Science, 136(5):jcs260574. doi: 10.1242/jcs.260574.

Pucilowska, J., Vithayathil, J., Tavares, E. J., Kelly, C., Karlo, J. C., and Landreth, G. E. (2015). The 16p11.2 deletion mouse model of autism exhibits altered cortical progenitor proliferation and brain cytoarchitecture linked to the ERK MAPK pathway. The Journal of Neuroscience: The Official Journal of the Society for Neuroscience, 35(7):3190–3200. doi:10.1523/JNEUROSCI.4864-13.2015. PMID: 25698753 PMCID: PMC6605601.

Rodal, A. A., Sokolova, O., Robins, D. B., Daugherty, K. M., Hippenmeyer, S., Riezman, H., Grigorieff, N., and Goode, B. L. (2005). Conformational changes in the Arp2/3 complex leading to actin nucleation. Nature Structural & Molecular Biology, 12(1):26–31. doi:10.1038/nsmb870.

Serafini, T., Colamarino, S. A., Leonardo, E. D., Wang, H., Beddington, R., Skarnes, W. C., and Tessier-Lavigne, M. (1996). Netrin-1 Is Required for Commissural Axon Guidance in the Developing Vertebrate Nervous System. Cell, 87(6):1001–1014. doi: 10.1016/S0092-8674(00)81795-X.

Short, K. M. and Cox, T. C. (2006). Subclassification of the RBCC/TRIM superfamily reveals a novel motif necessary for microtubule binding. The Journal of Biological Chemistry, 281(13):8970–8980. doi: 10.1074/jbc.M512755200. PMID: 16434393.

Sokolova, O. S., Chemeris, A., Guo, S., Alioto, S. L., Gandhi, M., Padrick, S., Pechnikova, E., David, V., Gautreau, A., and Goode, B. L. (2017). Structural Basis of Arp2/3 Complex Inhibition by GMF, Coronin, and Arpin. Journal of Molecular Biology, 429(2):237–248. doi: 10.1016/j.jmb.2016.11.030.

Spence, E. F. and Soderling, S. H. (2015). Actin Out: Regulation of the Synaptic Cytoskeleton. Journal of Biological Chemistry, 290(48):28613–28622. doi: 10.1074/jbc.R115.655118.

Spillane, M., Ketschek, A., Jones, S. L., Korobova, F., Marsick, B., Lanier, L., Svitkina, T., and Gallo, G. (2011). The actin nucleating Arp2/3 complex contributes to the formation of axonal filopodia and branches through the regulation of actin patch precursors to filopodia. Developmental neurobiology, 71(9):747–758. doi: 10.1002/dneu.20907. PMID: 21557512.

Spillane, M., Ketschek, A., Donnelly, C. J., Pacheco, A., Twiss, J. L., and Gallo, G. (2012). Nerve growth factor-induced formation of axonal filopodia and collateral branches involves the intra-axonal synthesis of regulators of the actin-nucleating Arp2/3 complex. The Journal of neuroscience : the official journal of the Society for Neuroscience, 32(49):17671–17689. doi: 10.1523/JNEUROSCI.1079-12.2012. publisher: Society for Neuroscience.

Spoerl, Z., Stumpf, M., Noegel, A. A., and Hasse, A. (2002). Oligomerization, F-actin interaction, and membrane association of the ubiquitous mammalian coronin 3 are mediated by its carboxyl terminus. The Journal of Biological Chemistry, 277(50):48858–48867. doi: 10.1074/jbc.M205136200. PMID: 12377779.

Stirnimann, C. U., Petsalaki, E., Russell, R. B., and Müller, C. W. (2010). Wd40 proteins propel cellular networks. Trends in Biochemical Sciences, 35(10):565–574. doi: 10.1016/j.tibs.2010.04.003. PMID: 20451393.

Striepen, J. F. and Voeltz, G. K. (2022). Coronin 1c restricts endosomal branched actin to organize ER contact and endosome fission. The Journal of Cell Biology, 221(8):e202110089. doi: 10.1083/jcb.202110089. PMID: 35802042 PMCID: PMC9274145.

Suo, D., Park, J., Harrington, A. W., Zweifel, L. S., Mihalas, S., and Deppmann, C. D. (2014). Coronin-1 is a neurotrophin endosomal effector that is required for developmental competition for survival. Nature Neuroscience, 17(1):36–45. doi: 10.1038/nn.3593.

Suo, D., Park, J., Young, S., Makita, T., and Deppmann, C. D. (2015). Coronin-1 and Calcium Signaling Governs Sympathetic Final Target Innervation. Journal of Neuroscience, 35 (9):3893–3902. doi: 10.1523/JNEUROSCI.4402-14.2015. publisher: Society for Neuroscience section: Articles PMID: 25740518.

Svitkina, T. M. and Borisy, G. G. (1999). Arp2/3 Complex and Actin Depolymerizing Factor/Cofilin in Dendritic Organization and Treadmilling of Actin Filament Array in Lamellipodia. Journal of Cell Biology, 145(5):1009–1026. doi: 10.1083/jcb.145.5.1009.

Svitkina, T. M., Bulanova, E. A., Chaga, O. Y., Vignjevic, D. M., Kojima, S.-i., Vasiliev, J. M., and Borisy, G. G. (2003). Mechanism of filopodia initiation by reorganization of a dendritic network. Journal of Cell Biology, 160(3):409–421. doi: 10.1083/jcb.200210174.

Taylor, A. M., Menon, S., and Gupton, S. L. (2015). Passive microfluidic chamber for long-term imaging of axon guidance in response to soluble gradients. Lab on a chip, 15(13):2781–2789. doi: 10.1039/c5lc00503e.

Thion, M. S. and Garel, S. (2017). On place and time: microglia in embryonic and perinatal brain development. Current Opinion in Neurobiology, 47:121–130. doi: 10.1016/j.conb.2017.10.004.

Tocchini, C. and Ciosk, R. (2015). Trim-NHL proteins in development and disease. Seminars in Cell & Developmental Biology, 47-48:52–59. doi: 10.1016/j.semcdb.2015.10.017.

Tsujita, K., Itoh, T., Kondo, A., Oyama, M., Kozuka-Hata, H., Irino, Y., Hasegawa, J., and Takenawa, T. (2010). Proteome of acidic phospholipid-binding proteins: spatial and temporal regulation of Coronin 1a by phosphoinositides. The Journal of Biological Chemistry, 285(9):6781–6789. doi: 10.1074/jbc.M109.057018. PMID: 20032464 PMCID: PMC2825472.

Uetrecht, A. C. and Bear, J. E. (2006). Coronins: the return of the crown. Trends in cell biology, 16(8):421–426. doi: 10.1016/j.tcb.2006.06.002. PMID: 16806932 publisherplace: England.

Urbina, F. L., Menon, S., Goldfarb, D., Edwards, R., Ben Major, M., Brennwald, P., and Gupton, S. L. (2021). Trim67 regulates exocytic mode and neuronal morphogenesis via SNAP47. Cell Reports, 34(6):108743. doi: 10.1016/j.celrep.2021.108743.

Weiss, L. A., Shen, Y., Korn, J. M., Arking, D. E., Miller, D. T., Fossdal, R., Saemundsen, E., Stefansson, H., Ferreira, M. A. R., Green, T., Platt, O. S., Ruderfer, D. M., Walsh, C. A., Altshuler, D., Chakravarti, A., Tanzi, R. E., Stefansson, K., Santangelo, S. L., Gusella, J. F., Sklar, P., Wu, B.-L., and Daly, M. J. (2008). Association between Microdeletion and Microduplication at 16p11.2 and Autism. New England Journal of Medicine, 358(7):667– 675. doi: 10.1056/NEJMoa075974. publisher: Massachusetts Medical Society.

Winkle, C. C., McClain, L. M., Valtschanoff, J. G., Park, C. S., Maglione, C., and Gupton, S. L. (2014). A novel Netrin-1–sensitive mechanism promotes local SNARE-mediated exocytosis during axon branching. Journal of Cell Biology, 205(2):217–232. doi: 10.1083/jcb.201311003.

Winkle, C. C., Olsen, R. H. J., Kim, H., Moy, S. S., Song, J., and Gupton, S. L. (2016). Trim9 Deletion Alters the Morphogenesis of Developing and Adult-Born Hippocampal Neurons and Impairs Spatial Learning and Memory. Journal of Neuroscience, 36(18):4940–4958. doi: 10.1523/JNEUROSCI.3876-15.2016. publisher: Society for Neuroscience section: Articles PMID: 27147649.

Wolterink-Donselaar, I. G., Meerding, J. M., and Fernandes, C. (2009). A method for gender determination in newborn dark pigmented mice. Lab Animal, 38(1):35–38. doi: 10.1038/laban0109-35.

Wong, E. W., Glasgow, S. D., Trigiani, L. J., Chitsaz, D., Rymar, V., Sadikot, A., Ruthazer, E. S., Hamel, E., and Kennedy, T. E. (2019). Spatial memory formation requires netrin-1 expression by neurons in the adult mammalian brain. Learning & Memory, 26(3):77–83. doi: 10.1101/lm.049072.118. Company: Cold Spring Harbor Laboratory Press Distributor: Cold Spring Harbor Laboratory Press Institution: Cold Spring Harbor Laboratory Press Label: Cold Spring Harbor Laboratory Press publisher: Cold Spring Harbor Lab PMID: 30770464.

Wurm, J., Konttinen, H., Andressen, C., Malm, T., and Spittau, B. (2021). Microglia Development and Maturation and Its Implications for Induction of Microglia-Like Cells from Human iPSCs. International Journal of Molecular Sciences, 22(6):3088. doi:10.3390/ijms22063088. number: 6 publisher: Multidisciplinary Digital Publishing Institute.

Yadav, S., Oses-Prieto, J. A., Peters, C. J., Zhou, J., Pleasure, S. J., Burlingame, A. L., Jan, L. Y., and Jan, Y.-N. (2017). Taok2 Kinase Mediates PSD95 Stability and Dendritic Spine Maturation through Septin7 Phosphorylation. Neuron, 93(2):379–393. doi: 10.1016/j.neuron.2016.12.006. PMID: 28065648 PMCID: PMC5267388.

Yung, A. R., Nishitani, A. M., and Goodrich, L. V. (2015). Phenotypic analysis of mice completely lacking netrin 1. Development (Cambridge, England), 142(21):3686–3691. doi: 10.1242/dev.128942. PMID: 26395479 PMCID: PMC4647218.

Zhang, Y., Chen, K., Sloan, S. A., Bennett, M. L., Scholze, A. R., O’Keeffe, S., Phatnani, H. P., Guarnieri, P., Caneda, C., Ruderisch, N., Deng, S., Liddelow, S. A., Zhang, C., Daneman, R., Maniatis, T., Barres, B. A., and Wu, J. Q. (2014). An RNA-sequencing transcriptome and splicing database of glia, neurons, and vascular cells of the cerebral cortex. The Journal of Neuroscience: The Official Journal of the Society for Neuroscience, 34(36):11929–11947. doi: 10.1523/JNEUROSCI.1860-14.2014. PMID: 25186741 PM-CID: PMC4152602.

